## Supplementary material for "Ambulacrarian insulin-related peptides and their putative receptors suggest how insulin and similar peptides may have evolved from Insulin-like Growth Factor": Suppl.Data-Revised

by Jan A. Veenstra

### Content

Spreadsheet 1 contains contains protein sequences for various insulin-like peptides and their putative precursors, as well as their genbank accession numbers or the terms Artemis or Trinity. Artemis indicates that the sequence in question was derived from a genome assembly, Trinity that the sequence was obtained using the Trinity program on a collection of transcriptome SRAs. In a few instances a combination of Trinity and Artemis was used to obtain a sequence.

Spreadsheet 2 contains the expression of some of these genes for a more limited number of species (there is little sense if the only transcriptome SRAs available for a species are from whole animals of unknown sex and unknown physiological status, or when then there are only very few such SRAs). The data contain the SRA identifier, the number of spots of each SRA and then for each protein of interest two numbers, one in blue, which is number of half spots that contains coding sequence of the protein of interest. The second number in black is the blue number multiplied by 1,000,000 and divided by the number of total spots, which yields a relative number.

The content of the remainder of this document is listed below:

The SRAs used in this study. 2

Figure S1. Sequences alignment of ambulacrarian IGFs. 6

Figure S2. Sequence similarity tree of ambulacrarian IGFs. 12

Figure S3. Sequence alignment of ambulacrarian GSSs. 13

Figure S4. Phylogenetic tree of ambulacrarian GSSs. 14

Figure S5. Sequence alignment of dilp7 orthologs. 15

Figure S6. Phylogenetic tree of ambulacrarian dilp7 orthologs. 16

Figure S7. Sequence alignment of ambulacrarian octinsulins. 17

Figure S8. Phylogenetic tree of ambulacrarian octinsulins. 19

Figure S9. Sequence allignment of ambulacrarian multinsulins. 20

Figure S10. Sequence similarity tree of ambulacrarian multinsulins. 24

Figure S11. Phylogenetic tree of bursicon and GPA2/GPB5 receptors. 25

Figure S12. Sequence alignment of various gonadulin receptors. 26

Figure S13. Sequence alignment of dilp7 receptors. 28

Figure S14. Sequence alignment GRL101 receptors. 32

### SRAs used in the analysis:

***Anneissia japonica***
SRR9663012, SRR9663013, SRR9663014, SRR9663015, SRR9663016, SRR9663017, SRR9663018, SRR9663019, SRR9663020, SRR9663021, SRR9663022, SRR9663023, SRR9663024, SRR9663025, SRR9663026, SRR9663027, SRR9663028, SRR9663029, SRR9663030, SRR9663031, SRR9663032, SRR9663033, SRR9663034, SRR9663035, SRR9663036, SRR9663037, SRR9663038, SRR9663039, SRR9663040, SRR9663041, SRR9663042, SRR9897705, SRR9897706, SRR9897707, SRR9897708, SRR9897709, SRR9897710, SRR9897711, SRR9897712, SRR9897713, SRR9897714, SRR9897715, SRR9897716, SRR9897717, SRR9897718, SRR9897719, SRR9897720, SRR9897721 and SRR9897722.

***Antedon mediterranea***
SRR6650067.

***Apostichopus japonicus***
ERR1193930, ERR1193931, ERR1193932, SRR166825, SRR414926, SRR414927, SRR414929, SRR414930, SRR771602, SRR771603, SRR771604, SRR771605, SRR771606, SRR863579, SRR934650, SRR934651, SRR1002331, SRR1002332, SRR1002333, SRR1657913, SRR1657914, SRR1657915, SRR1657916, SRR1657917, SRR1657918, SRR1657919, SRR1657920, SRR1657921, SRR1657922, SRR1657923, SRR1657924, SRR1657925, SRR1663411, SRR1663428, SRR1663429, SRR1663431, SRR1663432, SRR1663433, SRR1663436, SRR1663437, SRR1663438, SRR1663439, SRR1663440, SRR2006096, SRR2006097, SRR2006098, SRR2006099, SRR2006100, SRR2006101, SRR2006102, SRR2006103, SRR2006104, SRR2010219, SRR2012319, SRR2012383, SRR2012397, SRR2012405, SRR2014140, SRR2014144, SRR2014145, SRR2014150, SRR2014151, SRR2014153, SRR2014166, SRR2034609, SRR2089767, SRR2128001, SRR2128002, SRR5083072, SRR5083073, SRR5083074, SRR5083075, SRR5083077, SRR5083078, SRR5083079, SRR5083080, SRR5083082, SRR5083083, SRR5083084, SRR5083085, SRR5083086, SRR5083087, SRR5083088, SRR5817185, SRR5817186, SRR5817187, SRR5817188, SRR5817189, SRR5817191, SRR5817192, SRR5817193, SRR5817194, SRR5817195, SRR5817196, SRR5817197, SRR5817198, SRR5817199, SRR5817200, SRR5817201, SRR5817202, SRR5817203, SRR5817204, SRR5817205, SRR5817287, SRR5817295, SRR5839517, SRR5839518, SRR5839536, SRR5839538, SRR5839540, SRR5839546, SRR5839650, SRR5839684, SRR5839694, SRR5839713, SRR5839720, SRR5839725, SRR5839728, SRR5839730, SRR5839731, SRR5839732, SRR5839733, SRR5839734, SRR5839798, SRR5839799, SRR5839800, SRR5839801, SRR5839814, SRR5839840, SRR5839889, SRR5839890, SRR5839891, SRR5839892, SRR5839893, SRR5839897, SRR5839899, SRR5839900, SRR5839901, SRR5839902, SRR5839903, SRR5839904, SRR5839949, SRR5839950, SRR5839951, SRR5839952, SRR5839953, SRR5839954, SRR5840021, SRR5840022, SRR5840023, SRR5840024, SRR5840025, SRR5840027, SRR6004440, SRR6004441, SRR6004442, SRR6004443, SRR6004444, SRR6004445, SRR6004446, SRR6004447, SRR6004448, SRR6075435, SRR6075436, SRR6075437, SRR6075438, SRR6378158, SRR6856564, SRR6856565, SRR6856566, SRR6856567, SRR6856568, SRR6856569, SRR8258074, SRR8258075, SRR8258076, SRR8258077, SRR8258078, SRR8258079, SRR8258080, SRR8258081, SRR8258082, SRR8279868, SRR8801559, SRR8801560, SRR8801561, SRR8801562, SRR8801563, SRR8801564, SRR8801565, SRR8801566, SRR8801567, SRR9670443, SRR9670444, SRR9670445, SRR9670446, SRR9670447, SRR9670448, SRR9670449, SRR9670450, SRR9670451, SRR9670452, SRR9670453, SRR9670454, SRR9670455, SRR9670456, SRR9670457, SRR9670458, SRR9670459, SRR9670460, SRR9670461, SRR9670462, SRR9670463, SRR9670464, SRR9670465, SRR9670466, SRR9670467, SRR9670468, SRR9670469, SRR9670470, SRR9670471, SRR9670472, SRR9670473, SRR9670474, SRR9670475, SRR9670476, SRR9670477, SRR9670478, SRR9670479, SRR9670480, SRR9670481, SRR9670482, SRR9670483, SRR9670484, SRR9670485, SRR9670486, SRR9670487, SRR9670488, SRR9670489, SRR9670490, SRR11793939, SRR11793942, SRR11793943, SRR11793944, SRR12505006, SRR12505011, SRR12505013, SRR12505016, SRR12505017, SRR12830751 and SRR12830752.

***Asterias rubens***

SRR3087891.

***Florometra serratissima***
SRR3097584.

***Holothuria scabra***
SRR9125585.

***Holothuria arguiensis***
SRR8889053, SRR8892978, SRR8893063, SRR8896868, SRR8896869, SRR8898365, SRR8898366, SRR8906597 and SRR8906598.

***Lytechinus variegatus***
SRR1139214, SRR1661401, SRR1661399, SRR1661397, SRR1661409, SRR1661406, SRR1661395, SRR1661363, SRR1661081, SRR1661075, SRR1661113, SRR1660833, SRR1661077, SRR1661079, SRR1660831, SRR1661112, SRR1661111, SRR1661090, SRR9673387, SRR9673388, SRR9673385, SRR9673386, SRR9673383, SRR9673384, SRR9673381, SRR9673382, SRR9673389, SRR9673390, SRR9673407, SRR9673408, SRR9673409, SRR9673410, SRR9673403, SRR9673404, SRR9673405, SRR9673406, SRR9673401, SRR9673402, SRR9673391, SRR9673392, SRR9673393, SRR9673394, SRR9673395, SRR9673396, SRR9673397, SRR9673398, SRR9673399, SRR9673400, SRR9673379, SRR9673380, SRR9673377, SRR9673378, SRR9673375 and SRR9673376.

***Mesocentrotus nudus***
SRR6438347, SRR6438348, SRR6438349, SRR6438350, SRR6438351 and SRR6438352.

***Oligometra serripinna***
SRR2845424.

***Ophioderma brevispina***
SRR10742213.

***Patiria miniata***
SRR1138704, SRR1138705, SRR1138706, SRR1138707, SRR1138708, SRR1139189, SRR1139190, SRR1139191, SRR1139193, SRR1139194, SRR1139195, SRR1139196, SRR1139197, SRR1139198, SRR1139199, SRR1139201, SRR1139214, SRR1139215, SRR1139455, SRR2454338, SRR5398132, SRR5398133, SRR5398134, SRR5398135, SRR5398136, SRR5398137, SRR5398138, SRR5398139, SRR5398140, SRR5398141, SRR5398142, SRR5398143, SRR5398144, SRR5398145, SRR5398146, SRR5398147, SRR5398148, SRR5398149, SRR5986254, SRR6054712, SRR8580044, SRR8580045, SRR8580046, SRR8580047, SRR8580048, SRR8580049, SRR8580050, SRR8580051, SRR8580052, SRR8580053, SRR8580054, SRR8580055, SRR8580056, SRR8580057, SRR8580058, SRR8580059, SRR8580060, SRR8580061, SRR8580062, SRR8580063, SRR8580064, SRR8580065, SRR8580066, SRR8926376, SRR8926377, SRR12983012, SRR12983013, SRR12983014, SRR12983015, SRR12983016, SRR12983017, SRR12983018, SRR12983019, SRR12983020, SRR12983021, SRR12983022, SRR12983023, SRR12983024, SRR12983025, SRR12983026, SRR12983027, SRR12983028, SRR12983029, SRR12983030, SRR12983031, SRR12983032, SRR12983033, SRR12983034, SRR12983035 and SRR12983036.

***Patiria pectinifera***
SRR1139200, SRR1141045, SRR5229423, SRR5229424, SRR5229425, SRR5229426, SRR5229427, SRR5229428, SRR8627925 and SRR8627926.

***Pisaster ochraceus***

SRR1139197, SRR2846074, SRR10982207, SRR10982208, SRR10982209, SRR10982210, SRR10982211, SRR10982212, SRR10982213, SRR10982214, SRR10982215, SRR10982216, SRR10982217, SRR10982218, SRR10982219, SRR10982220, SRR10982221, SRR10982222, SRR10982223, SRR10982224, SRR10982225, SRR10982226, SRR10982227, SRR10982228, SRR10982229, SRR10982230, SRR10982231, SRR10982232, SRR10982233, SRR10982234, SRR10982235, SRR10982236, SRR10982237, SRR10982238, SRR10982239, SRR10982240, SRR10982241, SRR10982242, SRR10982243, SRR10982244, SRR10982245, SRR10982246, SRR10982247, SRR10982248, SRR10982249, SRR10982250, SRR10982251, SRR10982252, SRR10982253, SRR10982254, SRR10982255, SRR10982256, SRR11293187 and SRR11293188.

***Ptychodera flava***
SRR1029584, SRR2033704, SRR2033705, SRR2033710, SRR2033711, SRR2033712, SRR2033714, SRR2033715, SRR2033716, SRR2033717, SRR2033718, SRR2033719, SRR2033720, SRR2033721, SRR2033722, SRR2033723, SRR2033724, SRR2033725, SRR2033726, SRR2033727, SRR2033728, SRR2033729, SRR2033730, SRR2033731, SRR2033732, SRR2033733, SRR2033734, SRR2033735, SRR2033736, SRR2033737, SRR2033738, SRR2033739, SRR2033740, SRR2033741, SRR2033742, SRR2033743, SRR2033744, SRR2033745, SRR2033746, SRR2079098, SRR2079099, SRR2079100, SRR2079101, SRR2079102, SRR2079103, SRR2079104, SRR2079105, SRR2079106, SRR2079107, SRR2079108, SRR2079109, SRR2079110, SRR2079111, SRR2079112, SRR2079113, SRR2079114, SRR2079115, SRR2079116, SRR2079117, SRR2079118, SRR2079119, SRR2079120, SRR2079121, SRR2079122, SRR2079123, SRR2079124, SRR2079125, SRR2079126, SRR2079127, SRR2079128, SRR2079129, SRR2079130, SRR2079131, SRR2079132, SRR2079133, SRR12736734, SRR12736735, SRR12736736, SRR12736737, SRR12736738, SRR12736739, SRR12736740, SRR12736741, SRR12736742, SRR12736743, SRR12736744, SRR12736745, SRR12736746, SRR12736747, SRR12736748, SRR12736749, SRR12736750, SRR12736751, SRR12736752, SRR12736753, SRR12736754, SRR12736755, SRR12736756, SRR12736757, SRR12736758, SRR12736759, SRR12736760, SRR12736761, SRR12736762, SRR12736763, SRR12736764, SRR12736765, SRR12736766, SRR12736767, SRR12736769, SRR12736770, SRR12736771, SRR12736772, SRR12736773, SRR12736774, SRR12736775, SRR12736776, SRR12736777, SRR12736778, SRR12736779, SRR12736780, SRR12736781, SRR12736782, SRR12736783 and SRR12736784.

***Saccoglossus kovalewskii***
SRR071700, SRR071701, SRR071702, SRR071703, SRR071704, SRR071705, SRR071706, SRR071707, SRR071708, SRR071709, SRR071710, SRR071711, SRR071712, SRR071713, SRR071714, SRR071715, SRR071716, SRR071717, SRR071718, SRR071719, SRR071720, SRR071721 and SRR071722.

***Schizocardium californicum***
SRR2921981 and SRR2922012.

***Strongylocentrotus purpuratus***
SRR409091, SRR409092, SRR409093, SRR409094, SRR409095, SRR409096, SRR505583, SRR505584, SRR505585, SRR505586, SRR505587, SRR505588, SRR505589, SRR531843, SRR531853, SRR531860, SRR531948, SRR531949, SRR531950, SRR531951, SRR531952, SRR531953, SRR531954, SRR531955, SRR531956, SRR531957, SRR531958, SRR531964, SRR531996, SRR532046, SRR532055, SRR532074, SRR532121, SRR532143, SRR532151, SRR533746, SRR1012313, SRR1012339, SRR1012340, SRR1012342, SRR1012401, SRR1012403, SRR1041572, SRR1041901, SRR1042009, SRR1043060, SRR1043069, SRR1139792, SRR1765910, SRR1765938, SRR1765978, SRR1765979, SRR1765980, SRR1765981, SRR1765982, SRR1765983, SRR1765984, SRR1765986, SRR1765988, SRR1765991, SRR2080791, SRR2080792, SRR2080793, SRR2080794, SRR2080795, SRR2080796, SRR2080797, SRR2080798, SRR2080799, SRR2080800, SRR2080801, SRR2080802, SRR2080803, SRR2080804, SRR2080805, SRR2080806, SRR2080807, SRR2080808, SRR2080809, SRR2080810, SRR2080811, SRR2080812, SRR2080813, SRR2080814, SRR2080815, SRR2080816, SRR2080817, SRR2080818, SRR2080819, SRR2080820, SRR2080821, SRR2080822, SRR2080823, SRR2080824, SRR2080825,

SRR2080826, SRR2080827, SRR2080828, SRR2080829, SRR2080830, SRR2080831, SRR2080832, SRR2080833, SRR2080834, SRR2080835, SRR2080836, SRR2080837, SRR2080838, SRR2080839, SRR2080840, SRR2080841, SRR2080842, SRR2080843, SRR2080844, SRR2080845, SRR2080846, SRR2080847, SRR2080848, SRR2080849, SRR2080850, SRR2080851, SRR2080852, SRR2080853, SRR2080854, SRR2080855, SRR2080856, SRR2080857, SRR2080858, SRR2080859, SRR2080860, SRR2080861, SRR2080862, SRR2080863, SRR2080864, SRR2080865, SRR2080866, SRR2080867, SRR2080868, SRR2080869, SRR2080870, SRR2080871, SRR2080872, SRR2080873, SRR2080874, SRR2080875, SRR2080876, SRR2080877, SRR2080878, SRR2080879, SRR2080880, SRR2080881, SRR2080882, SRR2080883, SRR2080884, SRR2080885, SRR2080886, SRR2080887, SRR2080888, SRR2080889, SRR2080890, SRR2080891, SRR2080892, SRR2080893, SRR2080894, SRR2080895, SRR2080896, SRR2080897, SRR2080898, SRR2080899, SRR2080900, SRR2080901, SRR2080902, SRR2080903, SRR2080904, SRR2080905, SRR2080906, SRR2080907, SRR2080908, SRR2080909, SRR2080910, SRR2080911, SRR2080912, SRR2080913, SRR2080914, SRR2080915, SRR2080916, SRR2080917, SRR2080918, SRR2080919, SRR2080920, SRR2080921, SRR2080922, SRR2080923, SRR2080924, SRR2080925, SRR2080926, SRR2080927, SRR2080928, SRR2080929, SRR2080930, SRR2080931, SRR2080932, SRR2080933, SRR2080934, SRR2080935, SRR2080936, SRR2080937, SRR2080938, SRR2080939, SRR2080940, SRR2080941, SRR2080942, SRR2080943, SRR2080944, SRR2080945, SRR2080946, SRR2080947, SRR2080948, SRR2080949, SRR2080950, SRR2080951, SRR2080952, SRR2080953, SRR2080954, SRR2080955, SRR3017856, SRR3017857, SRR3593587, SRR3593588, SRR3593589, SRR3593590, SRR3593591, SRR3593592, SRR3593593, SRR3593594, SRR3593595, SRR3593596, SRR3593597, SRR3593598, SRR3593599, SRR3593600, SRR3593601, SRR3593603, SRR3593604, SRR3593605, SRR3593607, SRR3593608, SRR3593610, SRR3593612, SRR3593613, SRR3593615, SRR3593616, SRR3593618, SRR3593619, SRR3593620, SRR8863027, SRR8863028, SRR8863029, SRR8863030, SRR8863031, SRR8863032, SRR8863033, SRR8863034, SRR8863035, SRR8863036, SRR8863037, SRR8863038, SRR8863039, SRR9595500, SRR9595501, SRR9595510, SRR9595511, SRR9595512, SRR9595513, SRR9595514, SRR9595515, SRR9595516, SRR9595517, SRR9595518, SRR9595519, SRR9693264, SRR9693265, SRR9693266, SRR10002625, SRR10002626, SRR10002627, SRR10002628, SRR10002629, SRR10002630, SRR10002631, SRR10002632, SRR10002633, SRR10002634, SRR10002635, SRR10002636, SRR10002637, SRR10002638, SRR10002639, SRR10002640, SRR10002641, SRR10002642, SRR10002643, SRR10002644, SRR10002645 and SRR10002646.

```
                      1
Anneissia             QPNHVRRYCG SELTSELERR CALKGGFNSP KKQESP---- ---------- ----------
Oligometra            -EPKYRYYCG TSLVQKLESL CAPR-GYNGP KPGTSV---- ---------- ----------
Antedon               -SPKYARFCG SDLASELQRR CRSQ-GYNGP QQNSPL---- ---------- ----------
Florometra            -KPKYARYCG ANLASELERR CRTR-GYNGP EQKTHL---- ---------- ----------
Lytechinus-1          ---SFPLLCG QELVNAVAFV CDGR-GYYGQ PS-------- ---------- ----------
Hemicentrotus-1       ----FPLLCG QELVKAVAAV CNDR-GYYGQ PS-------- ---------- ----------
Strongylocentrotus-1  ---SFPLLCG QELVKAVAAV CNDR-GYYGQ PS-------- ---------- ----------
Apostichopus-1        ---SGQRYCG EALLEAMAYV CGDR-GYFGV TSGIRGRS-- ---------- ----------
Holothuria-1          ADGTGQKYCG EALLEALAFI CGDR-GYYGM TSGIHGRS-- ---------- ----------
Asterias-1            ---SPVQLCG RELTETLRSI CGDR-GYYSP GQSFS----- ---------- ----------
Pisaster-1            ---APVQLCG KELTDALKSI CGER-GYYSP GQSFS----- ---------- ----------
Acanthaster-1         ---AVIQVCG NDLLDALKSV CGDR-GFYSP PPGYS----- ---------- ----------
Patiria-1             ---SPIRVCG NELLDALMSV CGDR-GLYSP PEGYS----- ---------- ----------
Ophiothrix-1          ----WQRLCG TFLVDVVSQV CGER-GTYAD DSTHDLRKRS LDNFVDRGKM YSAGISKTSD
Lytechinus-2          -----YRLCG RELADALAVV CKGR-GYYID DSEI------ ---------- ----------
Hemicentrotus-2       -----FRLCG RELADALAVV CKGR-GYYID DSEI------ ---------- ----------
Strongylocentrotus-2  ----SFRLCG RELADALAVV CKGR-GYYID DSEI------ ---------- ----------
Apostichopus-2        ----AQRYCG TNLADALRIV CADR-GYYTQ ----KGAF-- ---------- ----------
Holothuria-2          ----SQRYCG SNLADALKII CADR-GYYDQ HHTVHGAF-- ---------- ----------
Asterias-2            ----WPKICG EQLVETVSLV CSTR-GFYSH RDS------- ---------- ----------
Pisaster-2            ----WQKICG EQLVETVSLV CSTR-GFYSH RDS------- ---------- ----------
Acanthaster-2         ----WQRICG EQLVETVSIV CNTR-GFYSH R E------- ---------- ----------
Patiria-2             ----WQKICG EQLVETVSVV CSTR-GFYSH RVE------- ---------- ----------
Ophiothrix-2          -----SYLCG SQIVQMMAVV CEGR-GYYYT EGSTGQICN- ---------- --------IV
Amphiura              -MAMWGRLCG KPLADMVALV CDGR---YYT GQ-------- ---------- ----------
Ophioderma            -MGMWGRLCG AQLANMVSFV CEGR-GYYYG EE-------- ---------- ----------
Saccoglossus          ----WDKLCG RTLVDVLALI CNGR-GYNSG SPKK------ ---------- ----------
Ptychodera            ----WDRLCG RSLADMLALV CHGR-GYYTD VSRQQ----- ---------- ---------K
Schizocardium         ----WDKLCG RTLADMLSLV CHGR-GYYTD VSKHR-IR-- ---------- ----------

                     61
Anneissia             -------YLW NQEKRIEATS ---------- -TERAKEINI VYECCHSACT EAFLDSFCLS
Oligometra            -------NLF --GKKNK FT ---------- -TTASNKKDI VAECCDTPCS EEMLARFCLS
Antedon               -------LFQ --DKKPQISA ---------- -SIRSKEKDI IDACCLSRCT EAFLDSFCLS
Florometra            -------LFQ --DKKVQNIG ---------- -TIRSNERDI IEACCRSPCT EEFLDSYCLS
Lytechinus-1          --KRSAGVFE LESRAKTFLK SGISRGELGR SKRRVRTGLI VTECCHNRCS VSHLESYCNP
Hemicentrotus-1       --KRSAGIFE LETRAKTFLK SGISRGETRR SKRGSRTGLI VTECCLNRCS VSHLESYCNP
Strongylocentrotus-1  --KRSAGIFE LETRAKTFLK SGISRGETRR SKRGARTGLI VTECCLNRCS VSHLESYCNP
Apostichopus-1        ---VSRSPFL TEERANSFLT NE-------R TRKTRRTGRI VTECCDNPCS QRNLESYCNV
Holothuria-1          ---VSRSPFL SEERANSFLT ND-------G TRNRRGTGRI VTECCENYCT TSVLESYCNF
Asterias-1            ---------- ---------- ---------- RRAPT-HDGI ATRCCQSLCE SSILETYCNL
Pisaster-1            ---------- ---------- ---------- RRAPT-HDGI ATKCCQSLCE SSILETYCNL
Acanthaster-1         ---------- ---------- ---------- RRTPATQTGI ATRCCISYCE TSVLEKYCNP
Patiria-1             ---------- ---------- ---------- RRTPATQSGI ATRCCISHCD TALLETYCNK
Ophiothrix-1          TNNAESNPFR GESEAKSFLG L--------- SRQKRRTGRI VHECCNNICN YRIIESYCNP
Lytechinus-2          --AQKDSPIV PRHVASSFLG SSS-ASAHSR QRRRVRSGQI VNECCDNECS NNIMESYCNK
Hemicentrotus-2       --SQKDSPIV PHHVASSFLG SSS-ASAHSR QRRRVRTGQI VNECCDKECS NNIMESYCNR
Strongylocentrotus-2  --AQKDSPIV PHHVASSFLG SSS-ASAHSR QRRRVRTGQI VNECCDKECS NNIMESYCNR
Apostichopus-2        ------PPVI PRQRASSFLT TDS---ENHH PRRLRRGTGI VTECCEKACD REVLETYCNP
Holothuria-2          ------QAII PRETASSFLS TNT---ENHP RRIRRRTGQI VTECCINHCT RGILETYCNP
Asterias-2            --KRDVEVFQ NERAAKSFLG SRIG------ SRQRRRTGRI ATECCDRICS FDIVESYCNP
Pisaster-2            --KRDVELFQ NERAAKSFLG SRIG------ SRQRRRTGRI ATECCERVCS FDIVESYCNP
Acanthaster-2         --KREAELFQ DERTAKSFLG THIG------ SRQRRRTGRI ATECCEKVCS YDIVESYCNP
Patiria-2             --KREAELFQ DERTAKSFLS SHIG------ SRQRRRTGRI ATECCEKVCS FDIVESYCNP
Ophiothrix-2          YSFLTDPPLL TERQAKTFT- KSG------- ---RTRTGKI VTECCDNHCD MQIIESYCAP
Amphiura              ----TDSPLL SERQAKSFTS NGG------- ---RKRTGKI VTECCDKHCN LQIIESYCAP
Ophioderma            ----SDSPIL PERQAKAFT- NGG------- ---RPRTGKI VTECCDNHCN LQIIESYCAP
Saccoglossus          -KVKRESPFR SGMEANDFFG NIS-SKEKRR QRRRSGSGKI VDECCHQACD YTTLESYCAP
Ptychodera            SRRARETVFA TQEEANGFFG VG---S--GR TKRRRGSGLI VVECCDKICD YSTIESYCAP
Schizocardium         --LPRETIFQ TQEDAHKFFG ASV-FG--ER TQRRRGSGLI VAECCEKSCD YATIESYCAP
```

Figure S1. Sequences alignment of ambulacrarian IGFs. In order to illustrate the sequence similarities in the putative D- and E- domains of these molecules Seaview[1] was used.

[1] Gouy M, Guindon S, Gascuel O. 2010. SeaView version 4: A multiplatform graphical user interface for sequence alignment and phylogenetic tree building. Mol Biol Evol. 27:221-4. doi: 10.1093/molbev/msp259.

```
                     121
Anneissia            SNKDEDTTVE SDTV------ ---ETTTITG KKRKPTRKPK N-PLKS---- ----------
Oligometra           PTPKDNTIVE SETM------ ---ETTTIVN TKKQRTKKP- K-PTRK---- ----------
Antedon              SNPEDDITVE SETI------ ---ETTTIVG SKKQLKTKKP K-TTRK---- ----------
Florometra           STPVDDTTAE IETV------ ---ETTTIIG SKKPRTKKPK S-DRRK---- ----------
Lytechinus-1         LPQDAV---- ---H------ ---DPE---- -----VH--- -----IRLEK SAEEDAD---
Hemicentrotus-1      LQPDAV---- ---H------ ---DAE---- -----VH--- -----IRLEK SAEEDADET-
Strongylocentrotus-1 LPPDAV---- ---H------ ---DAE---- -----VH--- -----IRLEK SAEEDAD---
Apostichopus-1       ATTQTTEIPT ELTT------ ---EGT---- -----TTEPA ASPRRNSR-- NIEADG--TA
Holothuria-1         AT----ELPT ELST------ ---ERT---- -----TTEPS ASPRRNSR-- HAGADI--TP
Asterias-1           PAPPSQTQPS TAAP------ ---TTT---- -----TKMAP LTE--DRRTK DVVVDY---S
Pisaster-1           PAASQTQT-- QASS------ ---AAP---- -----TKMAP LTE--DRRTK EVVVDY---S
Acanthaster-1        PSTSQSQTA- ---------- ---AAP---- -----PRITT TPD--ERRAN EIVVDE---T
Patiria-1            PATPQSQTA- ---------- ---AAP---- -----QRITS TSD--DRRAN EIVVDP---P
Ophiothrix-1         WPTTTT---- ---T------ ---ATT---- -----TQSPE PLPNEQ--EG GYLTDEIQMK
Lytechinus-2         RTPEVPS--- ESAI------ ---SEI---- ---------- --P-----SG DRTEDSTVRT
Hemicentrotus-2      RTPEVPP--- ESAI------ ---SEN---- ---------- --P-----SE EITEDSTLRT
Strongylocentrotus-2 RTPEVPP--- ESAI------ ---SEN---- ---------- --P-----SE EITEDSTLRT
Apostichopus-2       HVTPTLALAS --LV------ ---TSLM-TK SPTPPSSEPS SSSSSSSSSR N--EDKFPMT
Holothuria-2         RVTLEPTRPA VTSL------ ---SPLN-PR SQEPSGVQPQ ESPEEVHQEN S--RDNYPMG
Asterias-2           WPVAIE---- ---S------ ---RDP---- -----PL-SP VAPGRVREDK SADVDYMYNP
Pisaster-2           WPV-VE---- ---S------ ---RDV---- -----SL-SP VAPGRVREDK SADVDNMY-P
Acanthaster-2        WPVVED---- ---R------ ---DDP---- -----ML-AP VAPGRVRQDK SADADLLLRP
Patiria-2            WPVVED---- ---R------ ---DDP---- -----MLLAP VAPGRVRQDK SADADELLLM
Ophiothrix-2         LQEGQVKFTS RNLD------ -FVNEAN-NV VEEPSVRVPQ QVE-----EN KIVEETVPYV
Amphiura             LPEGQTRTQL RHWFLHEKEK ESANEEN-NT VEELPPPPPP R-------SE ERDEEQVPDV
Ophioderma           LPEGQTRGPE RHEFLHEKER ETITDQN-NR VEPPPLV-AE E-------AI VEEEETVPVI
Saccoglossus         LPEGVVA--- DDSL------ ---KRF---- ---------- --L-----SQ SFGNDFKDTA
Ptychodera           WPKDIDP--- AKKI------ ---EGF---- ---------- --K-----EG TWEEEDY---
Schizocardium        WPKDMDP--- ALRF------ ---AGF---- ---------- --K-----YG SWEDEDY---

                     181
Anneissia            -RK------- ----KPKGSS ---------- ---------E INSE-----Q SA----SNTE
Oligometra           SGR------- ----KNKDDP ---------- ---------D -TSE-----E SA----SNKD
Antedon              SGK------- ---------- ---------- ---------- ---------- SE----KNRD
Florometra           DGR------- ----KSKGGS ---------- ---------D -MSD-----D SA----SNRD
Lytechinus-1         -----EGRPQ --DG----PS ---------- --QL-----D ---------- ----------
Hemicentrotus-1      -----QGRPQ --DG----PS ---------- --QL-----D ---------- ----------
Strongylocentrotus-1 -----EGRPQ --DG----PS ---------- --QL-----D ---------- ----------
Apostichopus-1       AAA--GGGSG QGNGRGKGKG ---------- RHGK-----G NRRE-----Q DQ------TV
Holothuria-1         DGE--TPR-Q SNNGRNRGNP ---------N RHSE-----D N--------P ED------NI
Asterias-1           DQL---ATEG SQMSRVDGVL -----THDT- --------VT NRSKT-TTES NE--GSYDNE
Pisaster-1           DQL---VIEG SQMSRVDGAL -----TQDI- --------VT NRSKSATTES NE--GSYDNE
Acanthaster-1        GQT---GNTN SQMLRGGNAM -----GAGS- --------RA NGTKAPPTEV VD--GRSDDD
Patiria-1            SGH---TGDT APRGESAKAP -----GPSG- --------AA NATKTPVAEV DE--GMSVDE
Ophiothrix-1         HIT--RGQE- ---D----NS ---------- --V------D LLSE-----G SD------LR
Lytechinus-2         NDQSTERRTD DTDD----AT NLEVPSPDAD APDA-----T ATSDVERPLP EE----TTAV
Hemicentrotus-2      DGDSTEIRTD NTNP----AT NLEVPSPEAN APDA-----T ATSDVEQSRS DN----TTAV
Strongylocentrotus-2 DGESTEIRTD -TNP----AT NLEVPSPDAN TPDA-----T ATSDVEQPRS DN----TTAV
Apostichopus-2       DNA--LGEDY -EGPTNEGPL TSGEPTPTEN RIS------N GPRG-----P SS----NASS
Holothuria-2         DNV--VREDY -EPPTNGGAL TSGQPTPTEN RIS------S GLRE-----H SN----T-SI
Asterias-2           DVV--DVEEA --------NS ---------- --VI-----Q ---R-----E ED------LI
Pisaster-2           NAA--DVEEA --------NS ---------- --VI-----Q ---R-----E ED------LI
Acanthaster-2        DIA--EISE- --DK----SS ---------- --LL-----R QAAE-----K DE------PI
Patiria-2            QEA--AAGEQ FAEE----NS ---------- --LL-----R ---Q-----E EE------LI
Ophiothrix-2         SQA--IVGET TSD------G FDWDSDVSET HNILNGGGDM YINDQPRDDP S-----ENDE
Amphiura             SQD--LTF-- --S------D HDWEDSIESE --TLNEAIVM L-ADAPLTEQ ------NTEE
Ophioderma           NQE--LVD-T LSD------Q IDLEDSIESE TNLLNDALLM L-SDQVRENS EQRIGTENEE
Saccoglossus         NEDKLEI-VT VVRP----SH ---------- --DE-----M DGTE---TRI ED----NEHV
Ptychodera           -HRKYHP-ES VEQP----NA NPEEPTPEPT TTDL-----D KSLE---SES RD----NVDI
Schizocardium        -YRKYHP-EE FTQP----DL TTWIASTDSA NHDE-----H FTTA---TPS E---------
```

Figure S1-continued.

```
                     241
Anneissia            LPSQTEPTKD KNGR------ ----G----- ---------- ---DK---NK KDK--CNKKS
Oligometra           YETKTEKPRN KNGR------ ----EKNKD- ----KN---- --KDK---NK KNK--CNKKT
Antedon              DESQTENPRK DRQK------ ----EKNKN- ----K---DK NKKNR---EE RNK--CDKKT
Florometra           YESQTETPRN KNGR------ ----EKNKN- ----KEGRNK EDRNK---ED RNK--CDKKT
Lytechinus-1         ----TA--TG T-------M- -------LG- ----TDA--S ESRGR----- VRI-DAVEKV
Hemicentrotus-1      ----TA--TG T-------V- -------PE- ----TDT--S ETRGR----- VRI-DAVEKV
Strongylocentrotus-1 ----TA--TG T-------V- -------PE- ----TEM--S ETRGR----- VRI-DAVEKV
Apostichopus-1       DVTSAE--TE GNTE--PPR- -------PN- ----QQENNE DNRGT----- PDQ-SEE---
Holothuria-1         DETVTH--R- --TE------ -------TE- ----DQSRSN ENRGN----- RGK-GNR---
Asterias-1           EGAPY----D K-PDDSSPS- ERGESI--Q- ----DE---- -DNEV----- ----------
Pisaster-1           EGAPY----G K-TDDSSPS- ERGESI--P- ----DE---- -ENGV----- ----------
Acanthaster-1        DDAA------ ---GEINTS- ERVGSL--T- ----E---PD EETGR----- ----------
Patiria-1            EEAA------ ---GESNTS- ERGVSL--G- ----EDEDQE EQAQR----- ----------
Ophiothrix-1         DETPMP--TG RATGGRSPQ- -------LG- ----HDQSAD VDSGF----- HRF-GDIGAE
Lytechinus-2         -EKPKK--KD NGRGKNS-S- SESSTKKNR- ---------- TSKGMS-KEE -R------RR
Hemicentrotus-2      -EKPKK--KD NGKGKNS-S- LESSTKKNR- ---------- TSKGMS-KEE -R------RR
Strongylocentrotus-2 -EKPRK--KD NGKGKNS-S- LESSTKKNR- ---------- TSKGMS-KED -R------RR
Apostichopus-2       LELPTR--TS TATTNSSRIV TEGAHLAES- ----SDQSSL EDTDSE-SEA PTKGASHERT
Holothuria-2         EQIPA----- -RTGNNSRVI TEGVNLTDG- ----SEQSSL EDENSE-SET PTRSSDGERV
Asterias-2           DDIETQ--DQ EIEQDEEQN- -------MQ- ----T--LPE EDAED----- TDI-R-----
Pisaster-2           DAIETE--EH DLGQDEGNI- -------MQ- ----T--QSE VDVDG----- KDI-Q-----
Acanthaster-2        DDLDTL--EN EYADGGNVM- -------LQ- ----A---RE GVKEE----- G---AELGKE
Patiria-2            DALETL--EN KNAEGENVV- -------LQ- ----Q---PE EFEEV----- EEM-TEPETE
Ophiothrix-2         EHIPKE--KS LIKQ------ INVTES--Q- ----TDEILD EDTDSENSES KL---SIGAT
Amphiura             ELLHKK--DI KSG------- -TNTTM--A- ----TQDEAR EEDVIL-SES SD-----RTKG
Ophioderma           ELLQKE--KE ENAV------ TEMTEL--E- ----TQDELN EEEVSE-NRH RN---HLNKT
Saccoglossus         -TPPTK--PD VITETSSLI- LDDINVNKQI ISSNTSVEVK SKAGNT-K-- PK------RE
Ptychodera           -EESRS--KD KSAE----S- EESVTENDD- ------LKSE ETNDNQ-DEY SR------EE
Schizocardium        --ETTS---S ENGE----H- IKTVNKEDD- ------TQVT GTHELE-KDH KF------AE

                     301
Anneissia            --------RD K-NNKPCRRK SRK------D SRRKNR---- -----KKN-- ----------
Oligometra           L-------TD R-YGNPCVRK SKE-RKEKDR S-RKDR---- -----KR--- ----------
Antedon              L-------KD K-NGKPCVRK SKE-RKEKER S-RKTR---- -----KRN-- ----------
Florometra           L-------TD K-YGKPCTRK SKE-RKQKDR TTRKNR---- -----KRN-- ----------
Lytechinus-1         ISGRLVPTST MGSS--TPPS RKK-PRKD-K SERRKS---- --SREEKQ-- ----------
Hemicentrotus-1      LSERLIPTST TGSS--PSPS RKK-PRKD-K SERRNS---- --SREAKQ-- ----------
Strongylocentrotus-1 LSERLIPTST TGSS--PSPS RKK-PRKD-K SERRNS---- --SREAKQ-- ----------
Apostichopus-1       -----RPRD- -----RCRGS KKK-GKCR-N QDRVEE---- -EPTQDRIEE EPTTSR----
Holothuria-1         -----DNQN- -----NCRNS KKK-GSKK-K GNRKCRPGNE DSASEDGGNR RPPS------
Asterias-1           --------NK PEPNNIRDNS KERGRNRTHK GVSSER---- ----RANNSR RRGLSSERRG
Pisaster-1           --------NK QEPNNTRDNS KERGRNRTHK GVSSER---- ----RANSSR RRGLSSERRG
Acanthaster-1        --------DV ATNRPHKTHS KERSKNRTSK SER--R---- ----RRRTNR RRSS-SERRM
Patiria-1            --------DI PTSRPHKTHS KERSKNRTKG -EK--K---- ----RRKTGR RRGI-SERRT
Ophiothrix-1         V----EESDS TIDN----PS SDR------- ---------- ---------- ----------
Lytechinus-2         IASE----ER KASRERKKEL SRERRRRLKL QQRKDK---- --KK------ ----------
Hemicentrotus-2      IASD----ER RASRERKKEL SRERRKRLKM QQRKDK---- --KK------ ----------
Strongylocentrotus-2 IASD----ER RASRERKKEL SRERRKRLKL QQRKDK---- --KK------ ----------
Apostichopus-2       RKKQ----KK QRTRPPKKPK K---LSRE-R KERKKK---- --KEKERRER QRTISKKLRN
Holothuria-2         RTGPSRPSRT RRPRPPKPPK RKKLTSKE-R RERKKD---- --KKKTKNKN D---------
Asterias-2           -----EPEDV EESFPVPVPT KKR-RKVE-- GRRSKE---- --S-KNKG-- ----------
Pisaster-2           -----EPQD- EESFPVPVPT KKR-RKVE-- GRRSKE---- --G-KNKG-- ----------
Acanthaster-2        M----EEGGE KMPFPEVVPT KKR-RRVE-- GRRSRE---- --NSRDRN-- ----------
Patiria-2            R----EELEE EFPFPEVIPT KKR-RRVE-- GRRSRE---- --NSRDRS-- ----------
Ophiothrix-2         K-------GN SGRTKPTHRP TSR-HSRRKN SREKKK---- -NKTKTR-SR EKKGRK--RG
Amphiura             K-------GN RTHRPTHRPS SER-KSRRRN SKEKKR---- -NKSR--EKK NREGKK--KN
Ophioderma           K-------GN --SRTNKPSS RER-NSRRKN SRERKK---- -NKTKSGGSR ERKAKR--RN
Saccoglossus         KKDR----DN SSSKKRSHPK PKKSRKKQRL QRIRKK---- --AR------ ----------
Ptychodera           YLGR----DK GPFRKHKKAP TKKLFKEKKL SED-NK---- --KK------ ----------
Schizocardium        YWMN----ED KMSRKHKKA- -KRLMKER-- ------R--- --AR------ ----------
```

Figure S1-continued.

```
                       361
Anneissia              -----KKN-- ---------- ---------- -------KPT ---------- RRPKRWDSET
Oligometra             -----NKN-- ---------- ---------- -------KPT ---------- RRPKRWDSET
Antedon                -----KKN-- ---------- ---------- -------KPT ---------- RRPKRWDSET
Florometra             -----KKN-- ---------- ---------- -------KPT ---------- RRPKRWDSET
Lytechinus-1           ---A-RREER RRNRE-RSSG GKSRSGRRKN K------D-- -ND-----RS SRAKRQGLTL
Hemicentrotus-1        ---A-RREER RRNRE-RSSG GRSRSGRRKD K------D-- -ND-----RA SRAKRHGLNM
Strongylocentrotus-1   ---A-RREER RRNRE-RSSG GRSRNGRRKD K------D-- -ND-----RA SRAKRHGLNL
Apostichopus-1         --EVSSTDGL -SRERGSGSG GGRNRG-N-- ---------- ---------- GKGKGKGSQR
Holothuria-1           --AATATEGA -DRSSGSRGG RGSNHQ-TGR G------S-- -DRGSSRSNG GKGKGNRNQD
Asterias-1             SSSSRREEKL RRRRQ----- --------RH RERELREQRK QSN--SKRKS KGDKKDHSV-
Pisaster-1             SSSSRREAKL RRRRQ----- --------RH RERELREQRK QSN--SKRKT KGDKKDHSA-
Acanthaster-1          LSSERKREDA TRKLR----- --------RK EQRLSRKQP- -HS--NKRKS KLE-KKGSE-
Patiria-1              LSIERKKDDA TRRRR----- ---------K EQRLLRKQQQ SNS--SKRRT KLE-KKGSE-
Ophiothrix-1           ---------- ---------- ---------- ---------- ---------- ----------
Lytechinus-2           ---KKRLESA ERNR------ -GTDHMGLSE DSTLLAREPL GID--VR--- ---KRFH---
Hemicentrotus-2        ---KKRLESA ERNR------ -GTDHMGLSE DSTLLAREPL GID--VR--- ---KRFH---
Strongylocentrotus-2   ---KKRLESA ERNR------ -GTDHMGLSE DSTLLAREPL GID--VR--- ---KRFH---
Apostichopus-2         EERAAGLNPT EQQEDGGGGG -GGGGGGGGG G------EPR NNSNSDQPRN NRGKVNESQR
Holothuria-2           --RGQRTNIK -KQKSGERGS ANPNPTRGGD R------EPT D----NEQSR NRGKVNESQR
Asterias-2             ---G-KSEGK NKKRSGSREG GRSSRRSRGK S------S-- -RS-----KK QRDGRERSKR
Pisaster-2             ---G-KSEGK NKRRSGSREG GRSSRRSRGK S------S-- -RS-----KK QRDGRERSKR
Acanthaster-2          ---G-KSEGK SKRKSGSREG GRSFRR-RGK N------S-- -RR-----KK GRDGRERSKR
Patiria-2              ---G-KSEGR SRKKSGSREG DRSSRR-RGK S------S-- -RR-----KK GRDGRERSKR
Ophiothrix-2           K-PKKNKNKH DRRRKGKRHG GD------RD DST----QPT ---------- ----------
Amphiura               K-SKNK---K NRRRKN---R HG------ED DFT----QPT ---------- ----------
Ophioderma             K-PKHK---K NRRRKP---K HE------QD DYT----QPT ---------- ----------
Saccoglossus           ---GTRKTKL RHVKVKSKST PI-QQI---E TTTTM--KPF FDV---D-TD IIFYRFVNKT
Ptychodera             ---KSRAKSK NSTKVKPT-- -YVSSMTTSD EETLT--RPQ GDE--ARRET DATRKTEDSS
Schizocardium          ---KSRTKKG NNTKSKSILG KLVENM---- ---------- --E--TESPT NWQRDGTDER

                       421
Anneissia              TTLSDFLFSQ RFLQRAYDSD ID----VDEI TTEFGVVPEE DISSSLSEEE LSSNQVKSDD
Oligometra             TTLSEF---- --LQDIYNSD FADLSNIDKI KTGFDTIPEG DPSSSVSQEE SSSNEVIPDA
Antedon                STLSDFLLSQ KLLQEIYDSD IDDRS-IDDI TTQFDALPED DLSSSLSQEE SSSNEVIQEA
Florometra             TTLSDFLLSQ RFLQGVYDSD IGDQSTLDEL APRFDAVQEG DTSYSLSQE- SSSNEVISEA
Lytechinus-1           WRN---MFSQ KALSN----- ---------- ---------- ---------- ----------
Hemicentrotus-1        WRN---MFSD KGFSN----- ---------- ---------- ---------- ----------
Strongylocentrotus-1   WRN---MFSD KFFSD----- ---------- ---------- ---------- ----------
Apostichopus-1         REQSEELTVE ------TTTG CASQDGQN-- ---CEESNAN AGGAS----- ----------
Holothuria-1           RNQTDEDAER NTSSE-NGHG SRQRGGGG-- ---EGEERRN NGGSSQRGGT SNSNTGTGNS
Asterias-1             -AATTPLAVQ ---------- ---------- ---E--RPLK NGGRNSTSGE HS-----SVN
Pisaster-1             -AVTTPLAVQ ---------- ---------- ---EPIHPLK NGGRNSTSGE LS-----SVN
Acanthaster-1          -SVTTPVAVQ ---------- ---------- ---TADHPFK HGAYNSTTGD LS-----LVN
Patiria-1              --VTTPVAIE ---------- ---------- ---TAEHPFK NGGYNSTSGD LS-----PVN
Ophiothrix-1           ---------- ---------- ---------- ---------- ---------- ----------
Lytechinus-2           ----HT---P RSSRE----- ---QAST--- ----ATHTSD DD-----PAT ------SRQE
Hemicentrotus-2        ----HT---P RSSRE----- ---QAST--- ----ATHALD DD-----SAT ------SRQE
Strongylocentrotus-2   ----HT---P RSSRE----- ---QAST--- ----ATHALD DD-----PAT ------SRQE
Apostichopus-2         HPN------- ---------- ---------- ---------- ---------- ----------
Holothuria-2           PAE------- ---------- ---------- ---------- ---------- ----------
Asterias-2             WEG---LDTS HPVKEPTARS V---LGR--- ---------- VDTRPFRNFL --YNRYTVDE
Pisaster-2             WEG---LDTS HPEKEPAARS V---FGG--- ---------- IDTRPFRNFL --YNRYTVDE
Acanthaster-2          WEA---LVTS HPPTKDIVDA LRSALRA--- ---------- AGRSPSRVA- ---GSFAPDE
Patiria-2              WEG---LVTS HPPTKEIVDA LRSALRA--- ---------- AGRTPSLFAP --ANNFLLDE
Ophiothrix-2           DEFITDIEVP HPRIQAVAGM PSSRSEPA-- ---NIDDPTD PGTRSERSFV ----------
Amphiura               DELITDIEVM HPQARVVSRS EPVH------ ---AVDDPTD PGSRSERSFL ----------
Ophioderma             DQLITDIEVL HPQVQVVNAE SPSRSEAG-- ---NLDEAAD AGSRSERSFL ----------
Saccoglossus           WRVIEE---P EIMV------ ----GGK--- ----SESSRE SDIDSSI--- -------SHN
Ptychodera             WWMVDN---R EFEY------ ----EGR--- ----DRHNKS KTRK-GGNAT ------KVKK
Schizocardium          WWTVED---P HFQT------ ----GET--- ----DENSSN DKKTNGNNGT ------KWAR
```

Figure S1-continued.

```
                     481
Anneissia            QRTEHLPA-- ---------- ---------- ---------- -DI------- EVQ-SFDNIM
Oligometra           QRTDPFRP-- ---------- ---------- ---------- -PYKHILVDP IDM-DSYDNM
Antedon              QRTEQFRP-- ---------- ---------- ---------- -PLKPVPADT SED-SYNNNM
Florometra           QRTEQFRP-- ---------- ---------- ---------- -PFRPIPADT SEE-SYNNNM
Lytechinus-1         -------T-- ---------- ---------- -QALENQHIL DSLNGIAPSS TIIDTFETET
Hemicentrotus-1      -------I-- ---------- ---------- -PGIQNQLNL HQVNGRAPSS TTIDTFQMQT
Strongylocentrotus-1 -------I-- ---------- ---------- -PGLENQPNL HPVNGRAPSS TTIDTFQMKS
Apostichopus-1       -RRPGKTGKG K-GNGRRDD- A--DISSSER NTGDIEEEGT VPTSGNRPS- --S-------
Holothuria-1         QRTPGKPKKP QKGSSRRNQN Q--GSEEADE NTGAVSDEQS ESRNRVTPTQ TNS-------
Asterias-1           GTETDTA--- ---------- --------G- -AGSPEVKKD DLITTITAVL SDMI------
Pisaster-1           GTETDTA--- ---------- ---------- --GSPEVKKD DLITTITAVL SDMI------
Acanthaster-1        VTDSDTA--- ---------- ---------- --PSSDTKKD GFFTTITAVL RDVI------
Patiria-1            VTDPDSP--- ---------- ---------- --PSSDTKKD GFFTTITAVL RDVI------
Ophiothrix-1         ---------- ---------- ---------- ---------- ---------- ----------
Lytechinus-2         RRRTQARL-- ---SSRERKL HRTTTATARE ---ESQRDRR NVLQRL---- ----------
Hemicentrotus-2      RRRTQSRP-- ---SSRERKA HRTTTATARE --EELQRERR NVMQRL---- ----------
Strongylocentrotus-2 RRRTQSRP-- ---SSRERKT HRTTTATARE --EEMQRERR NVMQRL---- ----------
Apostichopus-2       -----VAQ-- ---------- ---------- ------EM-- NPLDKITKRI LAS-------
Holothuria-2         -----RQQ-- ---------- ---------- ------NN-- EEIDKLKKKI TFI-------
Asterias-2           KRDTERE--- ---------- ---------- -----SYRAV APLTGY-NSH RGG-------
Pisaster-2           KRDTDKE--- ---------- ---------- -----SYRAV APLTGY-NAY RGA-------
Acanthaster-2        QRSPLGTP-- ---------- ---------- -AGLQHRSGL DP-SGKPRPS YRV-------
Patiria-2            QRSPQATP-- ---------- ---------- -TGLQHRSAL DLSSGKAGRS YRV-------
Ophiothrix-2         ---------- ---------- ---------- ---------- ---TTIHARL VGF-------
Amphiura             ---------- ---------- ---------- ---------- ---TTITAKL LDV-------
Ophioderma           ---------- ---------- ---------- ---------- ---TTITAKL LDV-------
Saccoglossus         SRESHEP--- ----SREIDT T-------DC --DSV----- ---------- ----------
Ptychodera           ARHSQQTD-- ---DSRSSAE G-------GN --SEEMDVWH KPLTRPRGQP RRV-------
Schizocardium        KRRKDKPE-- ---PAVSRGR Q-------RR --SAVLGHRN SIFKRLRDRS WMV-------

                     541
Anneissia            S--------- -----RVNSK K---KTAAIK LKELTNQFSR LRQMQDATNA PRE-------
Oligometra           FM-----NSD DLSVQNVKPR L---EDPEEK IKKLTQIFSA --IVHDSTRT P---------
Antedon              YV-----STG NTR-NEQPPL D---DDPVEK LKELRHIFAE --LEGDSTSS S---------
Florometra           FM-----NAG NSR-KGEPSL K---DDPVAK LKKIKTYLHI SGLMQDSTSS S---------
Lytechinus-1         LLQHEKSLQD GAISHTKQPS INADEDRNAI FSDLMTKLR- -TLVLDFTSD R---------
Hemicentrotus-1      SSPTQQ---R NGGENFEQSS INQEADKKMR FSALMTKLR- -TMVLKFPRD R---------
Strongylocentrotus-1 SIPIQDSPEG NGKENFEQSS INQEADKKMR FSALMTKLR- -TMVLKFPKD R---------
Apostichopus-1       ---RES-SQS NGKGGGKKKL SKAEKKRQR- ---------- -KLLRKQEKA DRKAERKTGK
Holothuria-1         ---NESSSRG GRGKGGGNKL SKAEKKRQR- ---------- -KLLRQQEKK RRKQERQTKS
Asterias-1           ---------- ---------- ---------- ---------- -GFQPD-NGN R---------
Pisaster-1           ---------- ---------- ---------- ---------- -GFQPD-NGN R---------
Acanthaster-1        ---------- ---------- ---------- ---------- -GFQHADDGN R---------
Patiria-1            ---------- ---------- ---------- ---------- -GFQQADDGN R---------
Ophiothrix-1         ---------- ---------- ---------- ---------- ---------- ----------
Lytechinus-2         ---------- ---------- ---------- --------T- -GIFL----- ----------
Hemicentrotus-2      ---------- ---------- ---------- --------T- -GLFL*---- ----------
Strongylocentrotus-2 ---------- ---------- ---------- --------T- -GLFL----- ----------
Apostichopus-2       ---------- ---------- ---------- -------IH- -DIEMNREAD ID--------
Holothuria-2         ---------- ---------- ---------- ----REALR- -EVIIEPDSQ IG--------
Asterias-2           ---------- --SQ------ ----PDNHPT ----LAALY- -NLAVKLAKG LQH-------
Pisaster-2           ---------- --SQ------ ----PDNHPT ----LAALY- -HLAVRLAKG LQH-------
Acanthaster-2        ---------- --TQ------ ----KDNQST ----LEALY- -NLALRLSRP RRG*------
Patiria-2            ---------- --TQ------ ----KDNQST ----LEALY- -NLAVRLSRP RRG-------
Ophiothrix-2         ---------- ---------- ---------- --------L- -ESQQQNNNR RRR-------
Amphiura             ---------- ---------- ---------- --------I- -GLQQNSNHD RRR-------
Ophioderma           ---------- ---------- ---------- --------M- -GLQQNSNTN VR--------
Saccoglossus         ---------- ---------- ---------- ---------- ---------- ----------
Ptychodera           ---------- ---------- -E--VTA--- ---------- ---------- ----------
Schizocardium        ---------- ---------- -K--HEA--- ----RTHFS- -NLLLQAARR V---------
```

Figure S1-continued.

```
                    601
Anneissia           ---------- ---------- ---------- ---------- ---------- ----------
Oligometra          ---------- ---------- ---------- ---------- ---------- ----------
Antedon             ---------- ---------- ---------- ---------- ---------- ----------
Florometra          ---------- ---------- ---------- ---------- ---------- ----------
Lytechinus-1        ---------- ---------- ---------- ---------- ---------- ----------
Hemicentrotus-1     ---------- ---------- ---------- ---------- ---------- ----------
Strongylocentrotus-1 ---------- ---------- ---------- ---------- ---------- ----------
Apostichopus-1      SKKSKSKNKN RQTKGDRERR TTVRWNSNDE RSRRQVPRDS FSSVNGKYNS GHPYRRVYSM
Holothuria-1        SKK---GKKG SRTQGERERR TTVRWNDGEE RSRRHVPREQ FSPNNPRTTY HTDSR-TSLF
Asterias-1          ---------- ---------- ---------- ---------- ---------- ----------
Pisaster-1          ---------- ---------- ---------- ---------- ---------- ----------
Acanthaster-1       ---------- ---------- ---------- ---------- ---------- ----------
Patiria-1           ---------- ---------- ---------- ---------- ---------- ----------
Ophiothrix-1        ---------- ---------- ---------- ---------- ---------- ----------
Lytechinus-2        ---------- ---------- ---------- ---------- ---------- ----------
Hemicentrotus-2     ---------- ---------- ---------- ---------- ---------- ----------
Strongylocentrotus-2 ---------- ---------- ---------- ---------- ---------- ----------
Apostichopus-2      ---------- ---------- ---------- ---------- ---------- ----------
Holothuria-2        ---------- ---------- ---------- ---------- ---------- ----------
Asterias-2          ---------- ---------- ---------- ---------- ---------- ----------
Pisaster-2          ---------- ---------- ---------- ---------- ---------- ----------
Acanthaster-2       ---------- ---------- ---------- ---------- ---------- ----------
Patiria-2           ---------- ---------- ---------- ---------- ---------- ----------
Ophiothrix-2        ---------- ---------- ---------- ---------- ---------- ----------
Amphiura            ---------- ---------- ---------- ---------- ---------- ----------
Ophioderma          ---------- ---------- ---------- ---------- ---------- ----------
Saccoglossus        ---------- ---------- ---------- ---------- ---------- ----------
Ptychodera          ---------- ---------- ---------- ---------- ---------- ----------
Schizocardium       ---------- ---------- ---------- ---------- ---------- ----------

                    661
Anneissia           ---------- ---------- ---------- --------
Oligometra          ---------- ---------- ---------- --------
Antedon             ---------- ---------- ---------- --------
Florometra          ---------- ---------- ---------- --------
Lytechinus-1        ---------- ---------- ---------- --------
Hemicentrotus-1     ---------- ---------- ---------- --------
Strongylocentrotus-1 ---------- ---------- ---------- --------
Apostichopus-1      DPRRTAA--H FRSKPKSVRQ IRRRPSSSSP RKVFIKAR
Holothuria-1        DQTRTSKTDQ HRGKHRKMSR NGKQLSRTRS KSSLLKPR
Asterias-1          ---------- ---------- ---------- --------
Pisaster-1          ---------- ---------- ---------- --------
Acanthaster-1       ---------- ---------- ---------- --------
Patiria-1           ---------- ---------- ---------- --------
Ophiothrix-1        ---------- ---------- ---------- --------
Lytechinus-2        ---------- ---------- ---------- --------
Hemicentrotus-2     ---------- ---------- ---------- --------
Strongylocentrotus-2 ---------- ---------- ---------- --------
Apostichopus-2      ---------- ---------- ---------- --------
Holothuria-2        ---------- ---------- ---------- --------
Asterias-2          ---------- ---------- ---------- --------
Pisaster-2          ---------- ---------- ---------- --------
Acanthaster-2       ---------- ---------- ---------- --------
Patiria-2           ---------- ---------- ---------- --------
Ophiothrix-2        ---------- ---------- ---------- --------
Amphiura            ---------- ---------- ---------- --------
Ophioderma          ---------- ---------- ---------- --------
Saccoglossus        ---------- ---------- ---------- --------
Ptychodera          ---------- ---------- ---------- --------
Schizocardium       ---------- ---------- ---------- --------
```

Figure S1-continued.

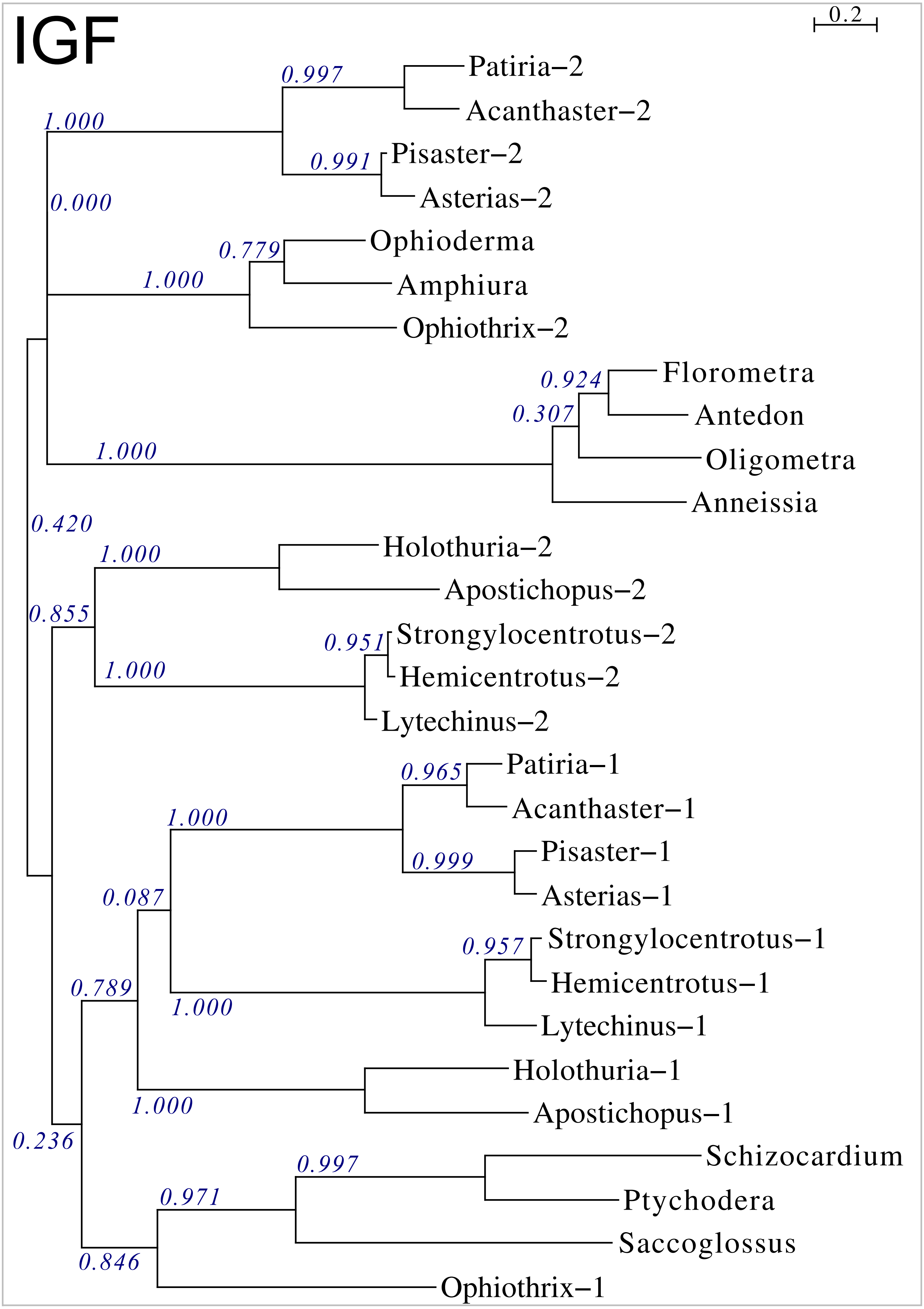


Figure S2. Sequence similarity tree of ambulacrarian IGFs.

```
Lytechinus            --QHLTRVRLCGLEFARAVYNHCSRASKRSDPGTISDIPLADRYLAR
Hemicentrotus         --QQGPRNRYCGLEFARAVFSQCSMANKRSDPGAVAESASAARYLAR
Strongylocentrotus    --QQGPRNRYCGLEFARAVFTQCSMANKRSDPGAVAESASAARYLAR
Apostichopus-1        -------IRLCGPDLSRAVYQICSHG-KRGYIPPTFNS--E------
Holothuria-1          -------VRLCGADLSRAVYRVCSHG-KRGYPMADL----E------
Apostichopus-2        ----WSHQRLCGPDLVHALSLVCGERG---YFGGSRLVER---D-VQ
Holothuria-2          -------HKLCGPALVDALAVVCRGRG---YYGGTVSRTK---REAS
Asterias-1            ------AEKYCDEDFHMAVYRTCTEH-KRS-GRSAFSL--N------
Pisaster-1            ------AEKYCDEDFHMAVYRTCAEH-KRS-GRSTYSL--N------
Acanthaster1          -------EKFCDNDFHLAVYQTCSTH-KRGDGEPVLSL--K------
Patiria-1             -------EKYCDDDFHMAVFRTCAVS-KRS--QPGMSL--S------
Asterias-2            RSDHASVKHFCGLEFSYAVVTACGEA-KRS-IRSAPFF---------
Pisaster-2            SNDHGRVKQYCGLEFSYAVVTACAEA-KRS-IRSAPFY---------
Acanthaster-2         ----DSSSKHCGSAFPQFVWTACSMA-KRS-NRSPRSL--D------
Patiria-2             ---TETTNRHCGAAFPDFVLAACSMA-KRS-IRSSPSL--H------
Ophiothrix-1          --DSARYQPLCGREFTRAVMEICATQVKRTEPLFQRFYNAN------
Ophiothrix-2          ----QDSYKSCGREFTRRVMEVCATHVKRTEHF--------------
Amphiura              -----DSAKYCGLAFSRAVMETCARQVKRTAPLWERLYTAS------
Ophioderma            -----DSATYCGVAFSMAVYETCSMQVKRTDPVRQRLYSAA------

Lytechinus            DTGYGQVQDTPFEWYDVAGQAEERLRP---------SL---SDIIFA
Hemicentrotus         DTGYEQAEDMPLEWYDIARQGAERLRP---------SL---TDIIFS
Strongylocentrotus    DTGYEQAEDMPLEWYDVARQGAERLRP---------SL---TDIIFS
Apostichopus-1        DDQLNQEFGTDL-----EEYLAETIKEYLKPNSLYDDVERELYPSLP
Holothuria-1          EEDFSQELDTEV-----DEFLAQALTGFLASRSFAADMESDRYYTLP
Apostichopus-2        DDGLDEEITTLV-------VGAERT-------SILECLK--AWSPFP
Holothuria-2          QDKLDV-------------------------------HA--GGSMTH
Asterias-1            D--F--------------------FRS-NSKRTAGSPRP--DDDFFL
Pisaster-1            D--LLSMD---------------SFRS-SPKRKAGSPLR--DDNSFL
Acanthaster1          D--VLTGS---------------RLRG-NIKRSFGSTLE--DEAFFA
Patiria-1             D--VLTMN---------------RFRGHNIKRSIDSTLE--DNAFFM
Asterias-2            D--MFPVF---------------KSPE-------RIPAD--FDDSSM
Pisaster-2            D--MFPVF---------------KSQE-------RIPAD--FDDSSM
Acanthaster-2         D--LLETF---------------KSAR-------HLDIS--Y--RTP
Patiria-2             D--LLQAF---------------KSDE-------YQANR--Y--TSP
Ophiothrix-1          ---LVKRSIDPAFWNNLLEANP--------------------DLMD-
Ophiothrix-2          ---MVKRSIDDEFWNDLMESGL--------------------GL---
Amphiura              ---RVKRFSDPEFWNAVLESDS--------------------DISM-
Ophioderma            ---RVRRFADPDFWEAVFNNDS--------------------NNVM-

Lytechinus            PRYRRSLFARPNTPMGRHCCVTGCTSQELASVC-------------
Hemicentrotus         SRFRRSIQNRRQLPMGQLCCVYGCTLVELASVC-------------
Strongylocentrotus    SRFRRSIHNRGQLPMGQLCCVYGCTLVELASVCT------------
Apostichopus-1        RG---FRRVTRTGGIARRCCSTGCSSSDIAKLC-------------
Holothuria-1          QR---FRRNA-RGGIARRCCASGCSSSDIAKLC-------------
Apostichopus-2        --------RRRTRGIVEECCFRRCTWENLESYCSKTTAYKKADNMI
Holothuria-2          --------VRFTRGIVEDCCFKGCTWSYLENYCLLS----------
Asterias-1            TM---QKRPETYVGMGSYCCLVGCTRDQLSQVC-------------
Pisaster-1            AL---QKRTETYVGMGSFCCLVGCTTEQLSQVC-------------
Acanthaster1          SR---LVKRSEYDGIASYCCIHGCTPSELAVVC-------------
Patiria-1             SG---LEKRSEYTGIASYCCLHGCTPSELSVVC-------------
Asterias-2            IH---VRKRQDYQGMATYCCTNGCTISQLTNSGIC-----------
Pisaster-2            IH---VRKRQDYQGMATYCCTNGCSISQLTNSGIC-----------
Acanthaster-2         IR---LSKRQDYDGMADYCCIIGCSTNELIASGIC-----------
Patiria-2             IH---LRKREEYMTIADYCCSVGCSPSDLVASGIC-----------
Ophiothrix-1          -----KRQSSAGVGMATHCCQSGCSQQEISMVC-------------
Ophiothrix-2          -----DKR--SETGMAEHCCQNGCTDQEISMVC-------------
Amphiura              -----DKRQPAGVGMATYCCNHGCTDTELSLVC-------------
Ophioderma            -----DKRQTTAMGMAQYCCSHGCTPQELSLVC-------------
```

Figure S3. Sequence alignment of ambulacrarian GSSs.

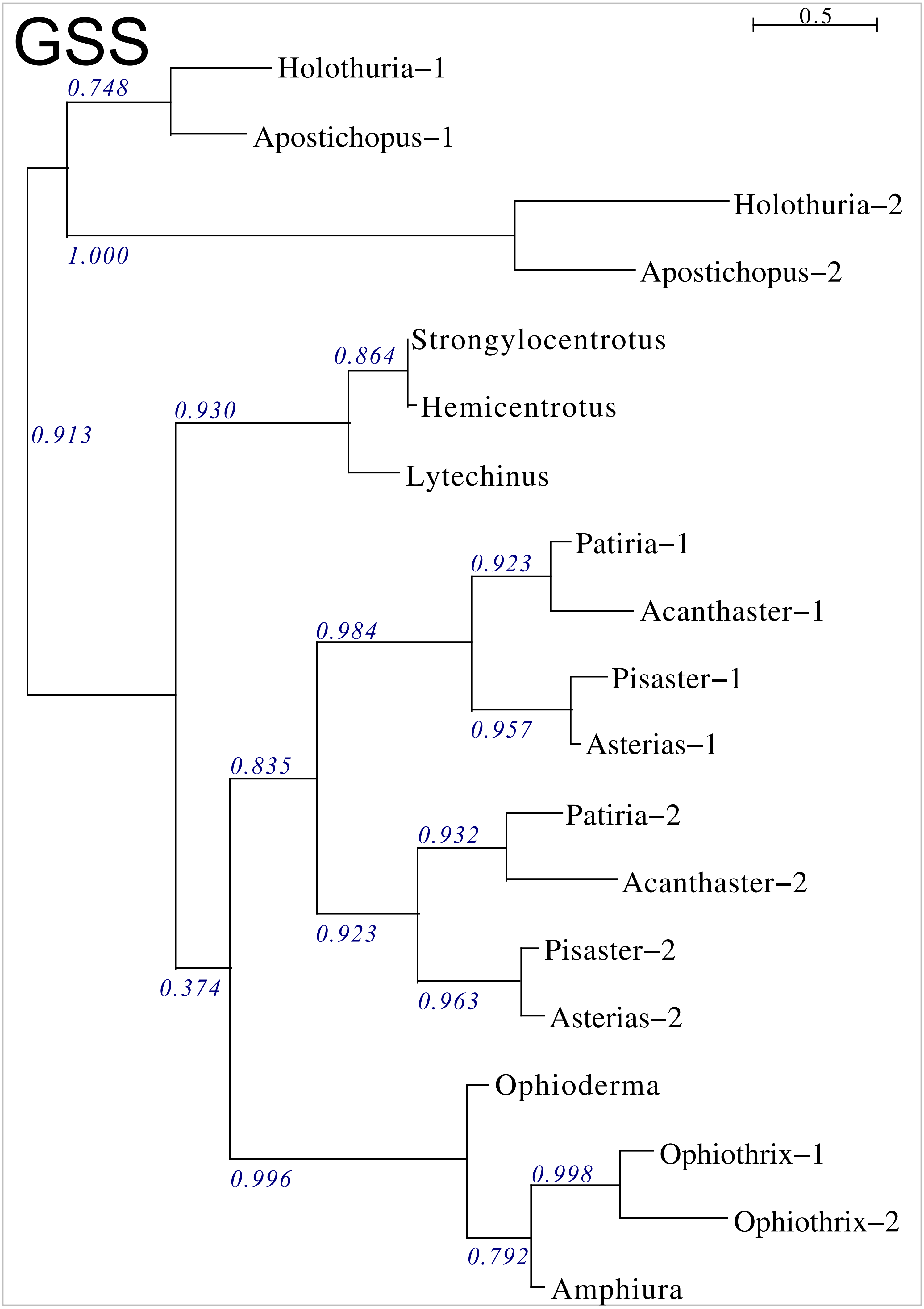


Figure S4. Phylogenetic tree of ambulacrarian GSSs.

```
Drosophila            ------LQHTEEGLEMLFRERSQSDWENVWHQETHS-R--C-RDKLVRQLYWA
Periplaneta           -------TKTENELEEMFKARSEEDWENAWHRERHT-R--C-QETLLRHLYWA
Stegodyphus           ------EVDVDPAWENIFKARSDEDWRSVWHTERHR-R--C-YHDLLVHMDWV
Oligometra            ---------------QDYSHRSRQEWARVWTVESMR-Q--C-HEDLREMVHIS
Anneissia             --------------LRDYSDRSHNDWARVWTVESMR-Q--C-HENLREMVHVS
Antedon               --------------LQDYSDRSHHDWARVWTVESMR-Q--C-HEDLREMVHIS
Florometra            --------------LQDYSDRSHYDWARVWTVESMR-Q--C-HEDLREMVHIS
Lytechinus            -DNLCNCQVIPQETMDEYELRTPEEWRENWLLDSLY-T--CVGPQLQRLGELA
Hemicentrotus         TEKFCNCMVIPELTMEDYEDRTPEEWRESWTMDTLR-T--CVGPQLQRVGELA
Strongylocentrotus    -EKFCNCMVLPELTMEDYEDRTPEEWRESWNMDTLR-T--CVGPQLQRVGELA
Apostichopus          -------------TLQELNSRTQPSWEQLWIVENV-PTVDCTV-DAVQLHIIA
Holothuria            -----DTTGLPETTIETLNAWTASEWVIAWASGSLVPVEGCTTLSVQQLFEYA
Asterias              --------AKPHLPVDQWRSRSKADWIRLWNTERHVNV--C-NEHLQPVFDAA
Pisaster              --------AKPHLPVDQWRSRSKADWIRLWNTERHVNV--C-NENLQPVFDAA
Acanthaster           ---------APHLPVEQWNSRSKADWVKLWNTERHVNT--C-NEALLPVWDVA
Patiria               --------AAPHLPVEQWKSRSKADWVKAWNTERHVDT--C-NEALLPVWDIA
Ophiothrix            -------------KITDYSSRTKADWQRLWLTESHQ-K--C-NEDLLPLWKIA
Amphiura              -------------RNTDYSSRTRADWQRLWVTESHQ-K--C-NEDLIPLWKIA
Ophioderma            -------------KTTDYSSRTTADWQRLWVTESHQ-K--C-NEDLIPLWKIA
Saccoglossus-1        LTVENEETITFSELNTMYGTRTLTDWQGKWTSETIH-A--C-GSNLYRISEYV
Ptychodera            ----AKDYDTVEDLILKYGTRTEDDWRNVWHTESHN-K--C-RNSLYSYVIFA
Schizocardium         ----ENAYETVESLKAKYSGRTFNDWRAAWNTETHN-K--C-RNSLYSYVIFA

Drosophila            CEKDIYRLTRRNKKRTGNDEAWIKKTTTEPDGSTWLHVNYANMFLRSRR----
Periplaneta           CEKDIYRLSRRNDYQQKENL-FLDE---GEPRYPFLSVVEARVFLRDRRRQR-
Stegodyphus           CQKDIYAVRRKKRST-----------------DPFLDDQKAHSFLGRRLKKSR
Oligometra            CRNDPRK-ITSKRN-------------------IFIPRSEATGFLSRFL----
Anneissia             CRNDPRK-ISSKRS-------------------IFIPRNEATGFLSRFL----
Antedon               CHNDPRK-ITSKRS-------------------IFIPRNEATGFLSRFL----
Florometra            CHNDPRK-ITSKRS-------------------IFIPRNEATGFLSRFL----
Lytechinus            CMNDPRKSVVVKRSN--------------SGREIFLPAKLAKAFLHYRHRHD-
Hemicentrotus         CINDPRKTIVVKRSN--------------SDRDLFLPAKLAKAFLQYRHRKD-
Strongylocentrotus    CINDPRKTIVVKRSN--------------SDRDLFLPAKLAKAFLHYRHRKD-
Apostichopus          CSNDVYKDHEGSRRRRF-------SDKRRRS--IFLNFSEANNFLA-KTKRT-
Holothuria            CQNDFYYDAEASTGL------GSGTGRRRRQSSIFLTHREASNFLLHRSKRFR
Asterias              CQNDVRK-IT-KRT-----------------GEEFVKEWAAKDFLIGS-----
Pisaster              CQYDVRK-IT-KRT-----------------GEEFVKEWTAKDFLIGS-----
Acanthaster           CQNDIRK-IT-KRM-----------------GREFLNEWTAKNFLAGS-----
Patiria               CQNDIRK-IS-KRT-----------------DREFVKEWNAKNFLAGS-----
Ophiothrix            CTYDIRK-IN-KRA------------------PEFVEDSEAKAFLIGP-----
Amphiura              CTYDIRK-IT-KRS------------------PEFIAEPDAKNFLLGP-----
Ophioderma            CTYDIRK-IN-KRS------------------PEFVAEPEAKNFLLGP-----
Saccoglossus-1        CYVDIHKSPDRKRTD-----------------DAFVDSAVAHDFLRGLMEKRT
Ptychodera            CMVDIYATGDKKRAE-----------------PPFVEHSIANSFLAGHRAEKR
Schizocardium         CMKDIHASGDRKRGQ-----------------FPFLDHEVADKFLSGSL-GKR

Drosophila            --SDGNTPSISNECCT-KAGCTWEEYAEY-CPSNKRRNHY-------------
Periplaneta           --RRGPGSSITEECCHNTAGCTWEEYAEY-CPANKRLRKFV------------
Stegodyphus           PHHTLVKRGIIDECCHGSAGCSWEEYAEY-CPANSRMRS--------------
Oligometra            --RTRRPSELHEDCCLDSRGCTWEEVAEIACINNRRRAHRPGSPIGR------
Anneissia             --RTRRPSELHEDCCLDSRGCTWEEVAEIACINNRRRMHRPGSPVGR------
Antedon               --RTRRPSELHEDCCLDSRGCTWEEVAEIACINNRRRMHRPGSPVGR------
Florometra            --RTRRPSELHEDCCLDSRGCTWEEVAEIACINNRRRIHRPGSPVGR------
Lytechinus            --SRRRRTGKDEECCLESNGCHLEELGEY-CTLHSRAYHVSGSQP--------
Hemicentrotus         --SRRRRVGKDEECCAEAQGCRWEELGEY-CTLHTRAYHQSGEQP--------
Strongylocentrotus    --SRRRRVGKDEECCAEAQGCRWEELGEY-CTLHTRAYHQSGEQP--------
Apostichopus          HSRVRRTTTFSTECCD--KLCIWEEVGEY-CWH-SRVYH--------------
Holothuria            RGRTRRSTTISAECCG-TQECKWEEIGEY-CVH-PREWHLP------------
Asterias              ----KRKRGLNEECCHEDTGCVWEEVAEY-CKKHGREKHKPGSTVAQGKQGR-
Pisaster              ----KRKRGLNEECCHEDTGCVWEEVAEY-CKQHGREKHKPGSPMVPGMQGR-
Acanthaster           ---KRRKRGLNEECCHEDLGCVWEEVAEY-CVMHGREKHEDGSPVRGKPGRRR
Patiria               ---RRRKRGLNEECCHEDLGCVWEEVAEY-CVMHGREKHEDGSPVRRRPGRRR
Ophiothrix            ---RRHKRGLNEECCHESKGCVWEEIGEY-CRMHSRASHVDGRIDSR------
Amphiura              ---IRHKRGLNEECCHEAKGCVWEEIGEY-CRMHSRASHVSGKVDAR------
Ophioderma            ---RRHKRGLNEECCHEAKGCVWEEIGEY-CRMHSRASHIDGRVDSR------
Saccoglossus-1        L-RRYRRTSATSECCADDGGCVWEELAEY-CTHQREVRTMDE-----------
Ptychodera            M-TRFRRASPTEECCSNAHGCNWEELAEY-CSHQRDDTRRR------------
Schizocardium         M-ARFRRANPSEECCSNRNGCTWEELAEY-CNHQRDDIDGR------------
```

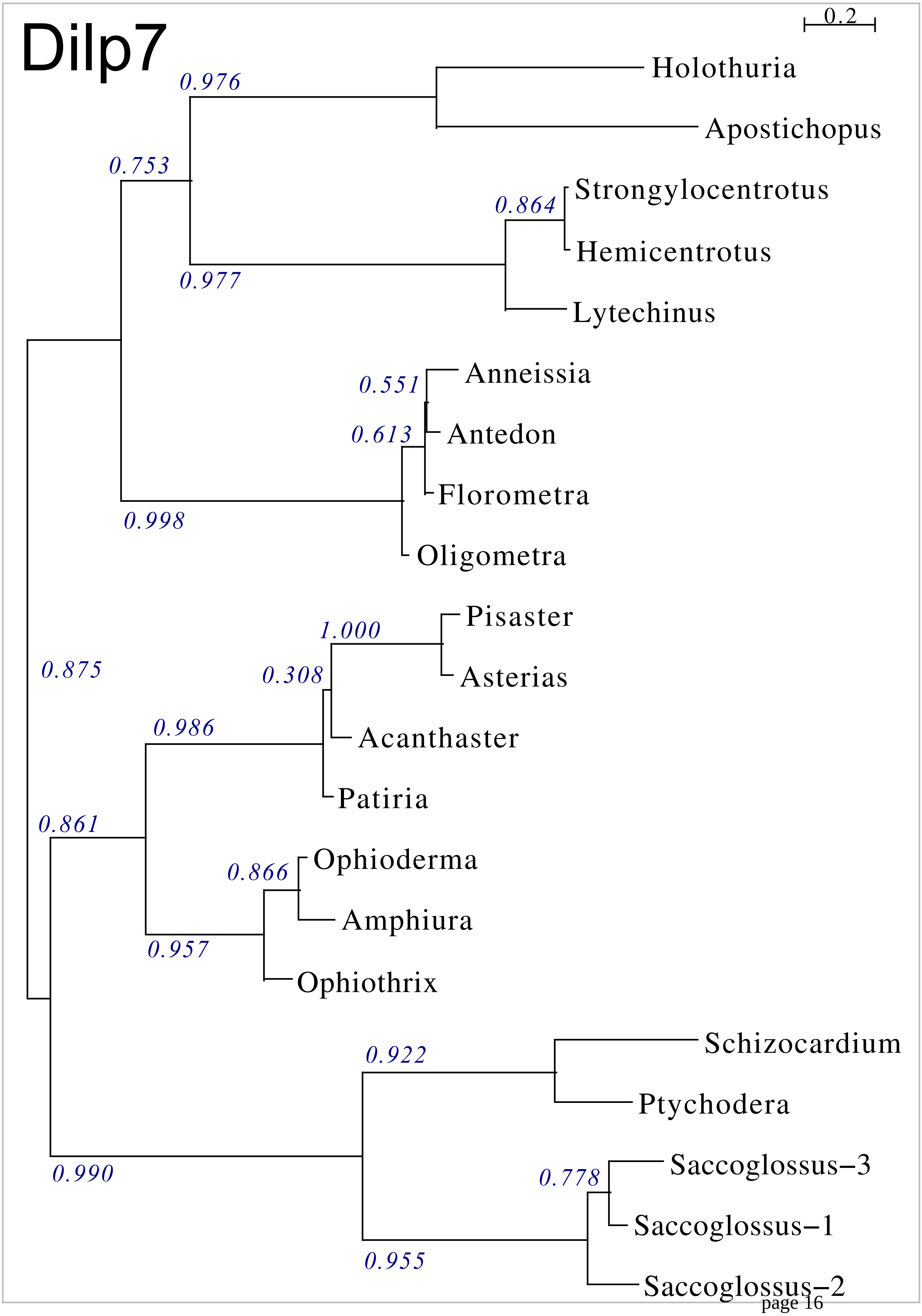




Figure S6. Phylogenetic tree of ambulacrarian dilp7 orthologs.

```
Anneissia            -------------------ARDWYC-GNAADTLKEFCQSCYASKRAHN--ALSLPSI
Oligometra           --------------------RDWYC-GNAATTLMDFCRSCYATKRAHT----SLPSS
Antedon              --------------------RDWYC-GSAANTLMNFCQSCYATKRSHS----SLPLL
Florometra           --------------------RDWYC-GNAATTLMDICQSCYATKRGYS----SLPSL
Lytechinus           --------------------QSWHC-GRAAQTIMSMCDSCYASYDKRS-------TS
Hemicentrotus        --------------------QSWHC-GRAAQTIMGMCNSCYASHDKRS-------IS
Strongylocentrotus   --------------------QSWHC-GRAAQTIMGMCNSCYASHDKRS-------IS
Apostichopus         ---------------------SWYC-GSAPETVRAICDGCYAGGIHTR--AFKRSSS
Holothuria           --------------------QSWHC-GSATETVRSVCNGCYAGGSHLSRNYLKRSHQ
Asterias             -------------------NNSWYC-SDVYSTLQSLCDSCYAGYAKRS-------DQ
Pisaster             -------------------NNSWYC-SDVYSTVQSLCDSCYAGYDKRS-------DQ
Acanthaster          --------------------DSWYC-SDVYSTVQSLCDSCYAGFDKRT-------NS
Patiria              ---------------------NWFC-SDVYSTMNSLCDGCYAGYDKRT-------NT
Amphiura             -----------------QRHERWYC-TPVFSMLQSMCGSCYAGIDKRS-------DN
Ophioderma           --------------------QRWYC-SPVFTMLQSMCGSCYAGIDKRS-------DD
Ophiothrix           --------------------NQWFC-SPVFTMLQSMCGSCYAGVDKRS-------DN
Saccoglossus-1       ------------------MSRNWHC-GRPVETMHEVCQGCYAGHVRPR-------NT
Ptychodera-1         ------------------LRREWHC-GRTVETMQGICRGCYAQPSERS-------T-
Schizocardium-1      ------------------FTKHWHC-GRIVDTMRAICDGCYASPTARD-------V-
Saccoglossus-2       RPSGSVD------------DVLTGC-KRKLLLVDQICAGCYAPPDIINNVNFDLTGK
Ptychodera-2         RPRNQGD------------D--VFC-SRTYSMVESVCDGCYATTQDSSPKSESAMNE
Schizocardium-2      RPNRPSE------------D--VRC-RKTIHMVLKLCNGCLAPIESEVENN--TISN
Saccoglossus-3       ---LPVDDTTTGVQVDVRDSRLWLC-GRLVEDLRALCRGCYAG---P----------
Ptychodera-3         -DLLPGNGDY---SNSAKGRRDLDC-THLVESMSLICRGCYATDQGV----------
Ptychodera-4         -----------------FLFNHRSCGTRLAETLSQICRGCYAHDDRP----------
Ptychodera-5         ----------------IRHRSGVSCGNRLRDSLASLCGGCYASDVR-----------
Ptychodera-6         -DLLPGNGDY---SNSANSRGNWHC-GRLVTTLSFICRGCYATGQ-V----------
Ptychodera-7         -DLLPGDGDY---SKSAKDRRDLDC-THLVDIMSFICRGCYATDQGV----------
Ptychodera-8         -DLLPGDGDY---SNSAKGRRDLDC-SLLIESMSFICRGCYATDQGV----------
Schizocardium-3      -------------QTSADRPRKWHCRKSVPEILSGVCRGCYAEPLQPP-------G-

Anneissia            KA-----KKDGMFLTKEGASGYLEAKRTRL--------FSS-LHLNHRQHETTNFVT
Oligometra           K------RTDGMFLTKERASGYLEAKRAKL--------FSSAVNLDRRQHAKTNFVT
Antedon              KS-----KSDEMFLTKERASGYLEAKRSRL--------FSS-VNLDRRQHGQTNFVT
Florometra           KS-----RTDGMFLTKERASGYLETKRSRL--------FSS-VNLDRRQHGQTNFVM
Lytechinus           KSSY---TPAKPFLHKRNAVHFLKSATREIENRPSIGDTAV-MSVERRSTGHRGFIH
Hemicentrotus        KSSY---TPAKPFLHKRNAVHFLKSTKREIESRPSIGDTAIEVAVERRSTGNRGFIH
Strongylocentrotus   KPSY---TPAKPFLHKRNAVHFLRTTKREIESRPSMGDTAIEVAVERRSTGNRGFIH
Apostichopus         DIIS---LYKDPFLKKSNALNFLLPRSHTP-------S-----SLIKRGIRRSGFIG
Holothuria           EKIP---LFKEPFLEKSYALNFLQAKSHAS-------DDPSPLMVRQRRSGRRGFID
Asterias             QTKT-----IDPFMDRKTAVDFFKRGTSR--------G-----------GARGGIID
Pisaster             QTKT-----IDPFMERKTAVDFFKRGASR--------G-----------GARRGIID
Acanthaster          ITRP---I-EEPFVERKNAVDFFKRGTAR--------G------------TRRGIVD
Patiria              ISRT---TDNEPFVERKTAVDFLKRGTAR--------G-----------STRRGIVD
Amphiura             VDRT-PQQSLDAFIQKEVAYSFIKRTSSG--------NS----FLRDSRSQHRGLID
Ophioderma           VDRT-QQQSLDAFIKKEVAHSFIKRTSAG--------SS----FLRNTR-QHRGLIE
Ophiothrix           SDTLSQKQSLDAFIQKEVAYSFIKRTSVG--------DT----FLRNARNTHHGLID
Saccoglossus-1       RS----VDGVQAFISRRDANMFTKGMSPD-------------V----KRAIDGGLIE
Ptychodera-1         -N----EAERQAFIGKEEASSFTKSVP---------------------TIVKRGLLE
Schizocardium-1      -G----SVKGLPFLKKHEASTFTRTRS---------------------SVQKRGIIE
Saccoglossus-2       --------------------------GPGY----------------EEEIVKNQIKE
Ptychodera-2         --------------------------SVQI----------------QDEDLREKIRD
Schizocardium-2      --------------------------ETPP----------------DQDVVTAKVRE
Saccoglossus-3       ------------DISKREASKFMQFNA-------------------HTIRQSRGIIE
Ptychodera-3         ------------TVNKRLASSFIPTTTTP-------------R-----NKIQRGIID
Ptychodera-4         ------------GISRREASSFLTTLRPQ-------------RAVARGSEERPNIVE
Ptychodera-5         -------------MTRRESSSFLQSK----------------------QRPLRGIVE
Ptychodera-6         ------------TVNKRLASSFIPTSTSP-------------R-----NKIQRGVID
Ptychodera-7         ------------TVDKRHASSFIPTTISP-------------S-----KKMQQDFIH
Ptychodera-8         ------------TVNKRLASSFIPTTTSP-------------R-----NKIQRGIID
Schizocardium-3      ---------KRDILNKQEASLFLRSP----------------------NGGDELIIN
```

Figure S7. Sequence alignment of ambulacrarian octinsulins.

```
Anneissia             ECCYNPCSSFEMIKYCCPTRQIELHNRNPNSSEDK----------------------
Oligometra            ECCYNPCSRFEMVKYCCTSRQIEFHESEGKK--------------------------
Antedon               ECCYNPCTRFEMIKYCCTSRQIEFNDTEGKK--------------------------
Florometra            ECCYNPCSRFEMIKYCCTSRQLEFNDTEGKK--------------------------
Lytechinus            ECCNKFCDPGEMILYCCEKRQIEWAHFHNLLKA------------------------
Hemicentrotus         ECCNKFCDPGEMVLYCCEKRQIEWAQFHNLLKA------------------------
Strongylocentrotus    ECCNKFCDPGEMVLYCCEKRQIEWAQFHNLLKA------------------------
Apostichopus          ECCEKNCEIREMVFYCCAEKQREYASFFPEIFRNRIRHT------------------
Holothuria            ECCRRSCDIFEMIFYCCAAQQQFYAEFFKLVKRS-----------------------
Asterias              ECCLKRCTTNELMLYCCEEKQREYFSFVGWLSRR-----------------------
Pisaster              ECCLKRCTTNEMMLYCCEEKQREYFSFVGWLARR-----------------------
Acanthaster           ECCHRQCAVSEMMLYCCEQKQREYYTFVGWLKRR-----------------------
Patiria               ECCHRPCSVNEMMLYCCEQKQREYYTFVGWLKRR-----------------------
Amphiura              ECCTSQCEAGEMILYCCQERQREWHSVRGFFNK------------------------
Ophioderma            ECCTQQCEIGEMVLYCCQERQREWHSMMGFFNK------------------------
Ophiothrix            ECCTQQCDTGEMILYCCQERQREWHIMMGLYNK------------------------
Saccoglossus-1        ECCYSQCSLTHMITYCCAEVQNEFQVFINILGNTDESSENDGDDGEESSSVHED---
Ptychodera-1          DCCYRRCNLQKMMTYCCAERQRELNNFFSLLNQKDNGST------------------
Schizocardium-1       HCCNHHCSFTELLIYCCEERSEEFYSFIGLLRMDDEDTDASLEKGNGDVEAEDNTDL
Saccoglossus-2        ACCKEYCPLPKIIEFCCDERQQEFHQFMSSFASTEE---------------------
Ptychodera-2          KCCNRRCTIHKMMQFCCEARRNEFHKFLALMGNTDN---------------------
Schizocardium-2       VCCDNYCSLDKIIEFCCEDLQQEFRQFMSFVSNSK----------------------
Saccoglossus-3        DCCYHTCPTERKIQYCCFEVQAQYRLFMESAI-------------------------
Ptychodera-3          ECCRNRCSVERKIQYCCDEIQKEFAFFFSLFGS------------------------
Ptychodera-4          ECCRRYCTLSRRIQYCCYEVQLEYQIYYESRNEE-----------------------
Ptychodera-5          ECCRSFCSLERRIQYCCYNVQARFAIFKEMSKEMSKERGM-----------------
Ptychodera-6          ECCRGTCSLERKIQYCCYEIQKEFALFKYGYRS------------------------
Ptychodera-7          ECCHTYCSFERKIQYCCNAVQQEFALFKSGYRS------------------------
Ptychodera-8          ECCHSRCSLQRKIQYCCDEIQEELALFLSLFGS------------------------
Schizocardium-3       ECCLRTCTVIEKIHYCCREKQIELYIIIQSAPWLVDNQR------------------

Anneissia             -------------------------------------
Oligometra            -------------------------------------
Antedon               -------------------------------------
Florometra            -------------------------------------
Lytechinus            -------------------------------------
Hemicentrotus         -------------------------------------
Strongylocentrotus    -------------------------------------
Apostichopus          -------------------------------------
Holothuria            -------------------------------------
Asterias              -------------------------------------
Pisaster              -------------------------------------
Acanthaster           -------------------------------------
Patiria               -------------------------------------
Amphiura              -------------------------------------
Ophioderma            -------------------------------------
Ophiothrix            -------------------------------------
Saccoglossus-1        -------------------------------------
Ptychodera-1          -------------------------------------
Schizocardium-1       DQGDDDDNTQNNEGGSIDVVVERDVGDVLLKTDKSKP
Saccoglossus-2        -------------------------------------
Ptychodera-2          -------------------------------------
Schizocardium-2       -------------------------------------
Saccoglossus-3        -------------------------------------
Ptychodera-3          -------------------------------------
Ptychodera-4          -------------------------------------
Ptychodera-5          -------------------------------------
Ptychodera-6          -------------------------------------
Ptychodera-7          -------------------------------------
Ptychodera-8          -------------------------------------
Schizocardium-3       -------------------------------------
```

Figure S7-continued.

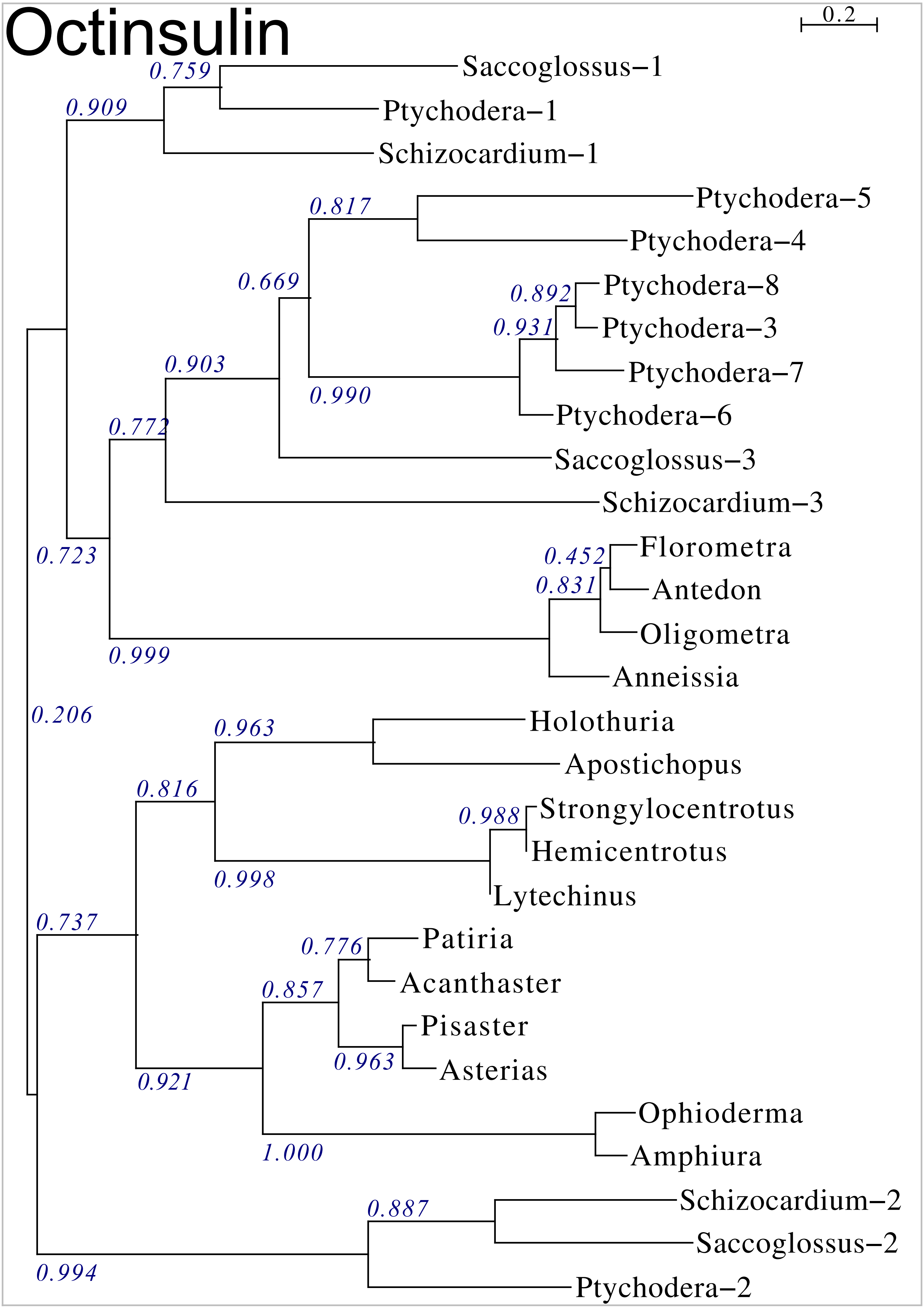


Figure S8. Phylogenetic tree of ambulacrarian octinsulins.

```
Amphiura-1      ---------------------------------------------------------
Amphiura-15     ------------------FD-----------LP------------------GREPGP
Amphiura-16     ----------------FLSDATSKKT-NEEALY------------------AQAMKR
Amphiura-2      ----------------YLTDAEH-------------------------------AER
Amphiura-3      ----------------YLTDAEH-------------------------------AER
Amphiura-7      YQDTDLRTLLDALEQSLRADSGDAPS-----------MKT-------------SKRA
Amphiura-6      IPDGYLRSLMDALQQIERADSGHAGV-DA------TAMKT-------------AERE
Amphiura-8      YQDTDLRTLLDELKQ--RADSRDAAA-----------IDT-------------NKRD
Amphiura-9      YQETDLRTLLEALKQ--REDSGDAPA-----------INT-------------NKKG
Amphiura-5      ------------------------------------------YP-----GALTRGTN
Amphiura-10     YPDTYLRTLLDALDQIVDSDATSKKT-NERAPDAYAMK-------------------
Amphiura-11     YPDTYLRTLLDALDQIVDSDATSKKT-NERAPDAYAMK-------------------
Amphiura-4      FPDNYLRTLLDALEKISDSDATGKRQ-MYADPDLARPPA------------------
Amphiura-14     FPDNYLRTLLDALEKIADSDATAKRG-LYDDEDLVRKPG------------------
Ophioderma-10   ---------------------DTETT-SSEEPDEWDEINP-TAIA--------ETKD
Ophioderma-3    -----------------------TKS-DNEEPDEWEVLHP-GE-------IREQSPD
Ophioderma-8    -----------------------TKS-ADEEPDEWEVLHP-GE-------PREPVSD
Amphiura-13     ------------------------KS-NSEEPDEHTT---------TTAPTTVGTDD
Amphiura-17     -----------------------TKS-IDEEPDELSVQGT-TTTGTTTAAVGRRDDD
Amphiura-18     -----------------------TKS-IDEEPDELSMQAT-T-----TVAARRRDDD
Ophioderma-1    ------------------DEIGAMKS-QNEEPDEHDSHIG-TT----QQ----PVIN
Ophioderma-2    -----------------------TKT-ANEEPDEWGLIPT-MP------PAPTDSLD
Ophioderma-4    -----------------------TNP-VNEEPDEGGLAP------------------
Ophioderma-7    -----------------------TTT-INEEPDEGGQAK------------------
Amphiura-12     -----------------------APNLFEDL-LRR--------DHVEQSGNITCM--
Acanthaster-5   ---------------------------VPRKPEEP--------SISAHCGMLLELRQ
Patiria-6       ---------------------------VPRKPGEP---------ISAHCGMLVELRQ
Asterias-4      -----------------------KLNLKPEETIDV--------TSSEHCGFVKEFRQ
Pisaster-2      -----------------------KLNMKPDEVLSV---------ESEHCSAVKEFRQ
Pisaster-5      ---------------------------------AQ--------QKSGHGPLLREVRQ
Asterias-1      -------------------------------------------ELRSQDGLMREVRQ
Pisaster-3      -------------------------------------------EPRPQDGLLREVRQ
Asterias-2      --------------------------------------------APNHNGLLREAKR
Pisaster-4      --------------------------------------------EPNHYGLLKETKR
Patiria-2       ----------------------DVPKPLA---DSE--------IDSNEDGEME----
Acanthaster-6   ----------------------DDPKPSESAAIAE--------DDSNKDGDIE----
Acanthaster-7   ----------------------DGPKLSES---------------TAIDSDIE----
Patiria-1       -----------------APPANDGIEVLTGGKIEE--------DDFEG----D-DFD
Acanthaster-2   ---------------------------------------------------LL-DLH
Acanthaster-1   ---------------------------------------------------HP-DAD
Acanthaster-8   ---------------------------------------------------HS-GAD
Acanthaster-9   ---------------------------------------------------HP-GAD
Patiria-5       ---------------------------------EP--------ADFNKDDKID-DLD
Patiria-8       -------------------------------------------------DKTD-DWH
Ophioderma-5    --------------NSLTRDADDAPVAMDDKPTSPRPIDIIKRSRHLRCAQFKDMAL
Ophioderma-6    ---------------------------------------AAVSPVHGSCVELK----
Ophioderma-9    ---------------------------------------AAFNSVPVSCADFQ----
Asterias-3      ------------------------------------------ELNVASCQQLADQVK
Pisaster-1      -----------------------------------------TELDEASCKQLADQVK
Acanthaster-3   ----------------------------------------DKTPSTASCEALAAQIS
Patiria-7       ----------------------------------------DKAPSAATCEALAATVK
Acanthaster-4   ---------------------SDTG--ADD---------DKVHIDDATCQALANQVK
Patiria-3       ---------------------DDAE--FKD---------GAIVIDDEVCEKLAAQVK
Patiria-4       ---------------------DDAE--FKD---------GAIVIDDEVCEKLAAQVK
```

Figure S9. Sequence allignment of ambulacrarian multinsulins.

```
Amphiura-1      ---------DWVGFRTICDPP-FTPSLYIEDFCGVT---VKR----------------
Amphiura-15     ASYV--ADNTWTGKTNICNYE-Q-QVVYHRELCGH----QKR----------------
Amphiura-16     YILN--PWSGWAGGKTECNTQ-TQESDF-DHACR------------------------
Amphiura-2      IDLA--KRKDW------CNNA---AHEF-SRLCN------------------------
Amphiura-3      IDLA--KRKDW------CNNA---AHEF-SRLCN------------------------
Amphiura-7      WEKE----FRSR---AECERQ-WK-LEL-DGHCISH----------------------
Amphiura-6      EGVK--RG----GVTPLCKSA-K--DRA-GKFCVYW----------------------
Amphiura-8      EGNE--KRFVSMSHSQACYSD-LYTYRK-PGICPTR----------------------
Amphiura-9      TDAR--D---------GCEAN-LN--VKVSQLC-------------------------
Amphiura-5      --DS--MNLRADFKLKLCGTE-LTLER--YYACGKR--DLK-----------------
Amphiura-10     ---R--GEYNWSGTKTLCGGT-LQTYK--GNICWGK--REAK----------------
Amphiura-11     ---R--GEYNWSGTKTLCGGT-LQTYK--GNICWGK--REAK----------------
Amphiura-4      --VF--DWQDWSGKKKICTAF-DQQQHF-NGVCQ------KRR---------------
Amphiura-14     --VF--DWEDWSGNKNICGRH-TQNQYY-NGVCQQYEGTGKRR---------------
Ophioderma-10   PVMM--RDKKWR-TSWFCGSD-IFRAK--KFICSPR--SYVRHIPY---------RKR
Ophioderma-3    IELD--HVKRWHSSVRICSS--KYRVQ--EAICDPR--NAKREVS-------------
Ophioderma-8    TELD--HVKRWHSSVRMCSS--MHRVK--ETICDPR--NAKREVS-------------
Amphiura-13     NAIL--MKRSWSGTKHLCGYQ-LNNER--SSVCGFK------RD--------------
Amphiura-17     DSST--LMKRWYGTVQLCGSH-LTKVK--SVICNPG--RPFKREE-------------
Amphiura-18     DSST--LMKRWNGYVQLCGSH-LTKVK--GVICNPG--APFKRDE-------------
Ophioderma-1    RSGT--MEKRWRGYVYICEPQ-LTRLK--NVICNPA--SVKRS---------------
Ophioderma-2    RSNM--LEKRWRGATKLCGST-LNTVQ--SVVCHPN--NYKRD---------------
Ophioderma-4    ------LGKRWSGSNKLCGEA-LSRAT--KNLCNPL--SAKRD---------------
Ophioderma-7    ------KGWSFTTGEKICGDD-LHTAT--TDVCFPD--SEKRD---------------
Amphiura-12     AYATMLEYEPNEEAYTWCDAN-LEHQR--EGICSCV--EYIKEHNKKRSLF-------
Acanthaster-5   MLDNPGLD-KRSPRNYWCGTT-LNNRK--SALCSCN--H------------------H
Patiria-6       MLDNPGLE-KRSPRNYWCGTA-LNNRK--STLCSCN--H------------------H
Asterias-4      LSDNPTLM-KRSPRNYWCSST-LERRK--EALCSCP--W------------------H
Pisaster-2      LADNPGVV-KRSPRNYWCNTA-LSQRK--TALCGCT--H------------------H
Pisaster-5      LFKGRNSQ-VRSETHYWCWEV-LEKNR--ESICNGP----------------PQPTQH
Asterias-1      LARNGRHF-RAAEDHIYCGVV-LEQNR--ESVCGVI----------------PGS---
Pisaster-3      LGRKPGHH-TRAEDHIYCGAV-LEQNR--EQVCGIV----------------PAS--R
Asterias-2      ILGRRGHQSRWDETHFYCDDA-LEQQR--EVICDPN--HTHAPVQPTPTGGPLPPTVS
Pisaster-4      VFRRRGFQPTNCEEAVEYCDA-LGQKR--EVLCQHN--HGTDAPRDSAVVG------A
Patiria-2       TKSGFSVA-WTGPQMWVCSSSTLAARK--HYVCHPP--GRHHHHKRE--------DE-
Acanthaster-6   AKGDFSVA-WSGPQMWVCRGR-LRRRR--RYACHPP--GRHHYKKRN--------DVQ
Acanthaster-7   AKGDFSVA-DNGPKLWVCQGN-VTNTL--KNVCTHT--GRPHSDTRD--------DVQ
Patiria-1       KNKNQYYS-RSGPRFRVCGTT-THSWS--SFVCHPS--GLI---------------HH
Acanthaster-2   AKEGRILV----RYPHYCNGI-LQIKY--NQLCHPQ--P-------------------
Acanthaster-1   AKHGGNLA----YRANYCGAT-LYEKV--RETCHAV--R-------------------
Acanthaster-8   TKEGGRVV----RYPHYCGAT-LQEKY--REICHAL--R-------------------
Acanthaster-9   TKEGGRVA----RYPHYCGAS-LQEKY--REICHAF--R-------------------
Patiria-5       EDDGQLET----RKPHYCGHR-LLRKY--AKICHHV--TLHRRPH-H--------LAA
Patiria-8       ADLG--QK----RSPHYCGRH-LGTKI--GQICGWH--L----------------EAW
Ophioderma-5    DHFRR-LVGKERVFAKYCSPT-PQTVM--DNYCQCD--VVPRSI-------------D
Ophioderma-6    ------KARELGAKHRFCGEE-LTYVR--NSYCSCR--VIPRDM-------------N
Ophioderma-9    ------SAQKRGNLHKLCGSE-LTAAV--DNYCQCK--VIPRDM-------------S
Asterias-3      EGKKRWD----GPSHKFCGET-LNEKR--YAYCTCG--LVPRKR-------------E
Pisaster-1      EGKKRWT----GSWHTFCGET-LNEKK--YAYCTCG--LVPRKR-------------E
Acanthaster-3   EGKKRWD----GPSHTFCGED-LNERR--NAYCNCQ--VVPRKR-------------E
Patiria-7       EGKKRWD----GPAHTFCGET-LTEKA--NAYCHCQ--VVPRKR-------------E
Acanthaster-4   ERGRRWPWSRTGPKHYFCQPT-LEKKK--EAYCNCK--VVPRKR-------------D
Patiria-3       EKS----ILGVGPKHWYCNPT-LERKK--EQYCTCS--VVPRER-------------D
Patiria-4       EKS----ILGVGPKHWYCNPT-LERKK--EQYCTCS--VVPRER-------------D
```

Figure S9-continued.

```
Amphiura-1      ------EYE-ATDPLGFLK-------MRGMVK---RLNP---NWRELMSEECCYESCT
Amphiura-15     ------EIE-AEDALGFLK-------MKLRGM---EKRG---GWQEYMNEECCIEWCT
Amphiura-16     -------LY-GRDALDFLK-------LRDVVK---RA--SFNDIKEWIHEECCLERCI
Amphiura-2      --------N-RRDALDFLN-------LRDVRN---KVHAFTNYIKEAVHEECCEESCI
Amphiura-3      --------N-RRDALDFLN-------LRDVRN---KLHAFTQYIKEGVYEECCDETCI
Amphiura-7      --PYRRDMP-DVDALAFLN-------KRG-------------KSLNGLYHECCDESCS
Amphiura-6      INRRDIEVS-MPDALGFL--------KTE-------------VQARDLNHECCVETCS
Amphiura-8      RSEDIQQLG-MQGALAFL--------KRE-------------MVKRDLFHECCVETCT
Amphiura-9      RREDIQQLD-TPDAFAFL--------KRE-------------LVQRDLYHECCVEQCD
Amphiura-5      --DALTFLE-DGA-----------------------------NRKRDLHEECCVETCS
Amphiura-10     --DALAFLE-TDDRQ----------------TDADP-LGIWALISTTLNEECCGEACY
Amphiura-11     --DALAFLE-TDDRQ----------------MDPLSTFISLALISTTLNEECCGEACY
Amphiura-4      --DIPQQLD-TPDALGFLQ-------AKRI-------AGFEKDVYESMTEECCLEGCT
Amphiura-14     --DIPQQLD-TPDALGFLQ-------AEK-----------RKDVFESINEECCAEQCS
Ophioderma-10   RSVGDATLI-KQDADMFLN-------RQKR-----------RKSSYSLYEECCGEFCS
Ophioderma-3    DNGVPGGVA-KETAKRFLK-------RQRR----------GEWGKGDLEEECCHEGCS
Ophioderma-8    DNGVPGGVA-KETAKRFLK-------RQRR----------GEWGKGDLEEECCHEGCS
Amphiura-13     AEDALDFLK-QDIV------------------------------KRSLTEECCHEGCS
Amphiura-17     SEAALAFLK-EERAKRFLQ-------ADQQLLHQQQLHQPSLRLWGGLTEECCNEHCS
Amphiura-18     GEAALAFLN-EERAKRFLQ-------PDQQ-----QLDQPNLRLWGGISEECCNEHCS
Ophioderma-1    DTAGLEFLT-EHQAKRFLM-------QNKR-----------IWGWGGLSEECCNEGCS
Ophioderma-2    --ETAALVG-EQQAKRFLR-------KR------------WWGGAGGLTEECCNEGCS
Ophioderma-4    --ETTAVME-ERRAKRFLI-----------------------EKRSSLNEECCNEGCA
Ophioderma-7    --A---IIQ-EKGAKRFLE-------NKRLW-------GWSWSTHGSLNEECCNEGCV
Amphiura-12     --KRMAEVT-EDVAKRFLESLLVLEVERAR------------RPKTSLQEECCNESCY
Acanthaster-5   SLLSDELIE-SKEANSFL--------KRGI------------EK-RSLTEECCHEGCY
Patiria-6       TFL-----K-SDEANNFL--------KRGV------------EK-RSLSEECCHEGCY
Asterias-4      TLRDDDFMEEKKDATNFL--------ERGV------------EK-RSLHEECCNEGCY
Pisaster-2      TLRDDDFMEEKKDATNFL--------ERGI------------EK-RSLSEECCHEGCY
Pisaster-5      YDSTTEFVG-AREAKGFL--------GKD---------------FTNLHEECCFEGCT
Asterias-1      --KRNLFVR-KEAASEFL--------ERG---------------VQGLAEECCGEGCT
Pisaster-3      LSKRNVFVG-KEAASDFL--------KRG---------------VQGLGEECCGEGCT
Asterias-2      PGHQDAFVG-KNEASRFL--------GRGV--------------VQGLIEECCFEGCH
Pisaster-4      IRSQDEFVK-RREASKFL--------GRGA--------------FQGLLEECCHEGCQ
Patiria-2       -DNSDAFLT-EDDANTFLA-------SPSK------------RKANIIHEECCHEGCF
Acanthaster-6   EESNPEFLD-EAEANFFLR-------R---------------HKRNHISEECCHEGCY
Acanthaster-7   EENNPEFLD-EADANFFLR-------Q---------------QNSKQISEECCHEGCY
Patiria-1       KRDNDEFLS-AGEANTFLM-------SEGV------------GK-RGMHEECCHEGCW
Acanthaster-2   EVFDQDFLD-SKEANTFLF-------KQRI------------GK-RSLIEECCFEGCT
Acanthaster-1   GVSNQEFLD-SKDASTFLF-------GRGI------------EK-RGLHHECCTEGCS
Acanthaster-8   GVSNQEFLD-SKDANTFLF-------GRGI------------GK-RNLNEECCSEGCD
Acanthaster-9   GVSNQEFLD-SKDANTFLF-------GRGI------------GK-RNLNEECCGEGCD
Patiria-5       NEAPSAFVE-SDEANSFLD-------NRGI------------SK-RGLHEECCHEGCS
Patiria-8       DDVHPAFVE-SDEANSFLA-------DRRI------------SK-RSLGEECCHEGCS
Ophioderma-5    --DKRAFVD-KSSAKSFLN--------HRA------------ST-RSLDEECCNEGCN
Ophioderma-6    P-------G-LEDAKAFLK-------MRSP------------ST-RSISEECCYETCY
Ophioderma-9    P-------G-LEDAKAFLK-------MRSP------------ST-RSISEECCYETCY
Asterias-3      -LELSEFLN-RGKANGFLS-------ARNV------------QK-RSLSEECCHEGCY
Pisaster-1      -LDLSEFLN-RGKANGFLS-------ARNL------------QK-RSLSEECCSEGCY
Acanthaster-3   -LDLSEFLP-SGKANAFLS-------GRRI------------TK-RSLSEECCHEGCY
Patiria-7       -LELSEFLT-PVKANSFLS-------GRNI------------AK-RSLTEECCHEGCY
Acanthaster-4   DGDQSEFLK-PEKAGSFLE--------SSI------------GK-RDFNHECCKEGCY
Patiria-3       -ADRAEFLE-PEEAGGFLN--------R--------------SK-RDLNHECCDEGCV
Patiria-4       -ADRAEFLE-PEEAGGFLN--------R--------------SK-RDLNHECCDEGCV
```

Figure S9-continued.

```
Amphiura-1      SEEIKELC-----------------
Amphiura-15     MEEVKELDC----------------
Amphiura-16     VEEVVELC-----------------
Amphiura-2      NEEVLEFC-----------------
Amphiura-3      DEEVLEFCP----------------
Amphiura-7      MEEFGEHCTQRPESQEGFNPFKLGK
Amphiura-6      DEEVKEHCVMTK-------------
Amphiura-8      TEEIQETC-----------------
Amphiura-9      AEEIEETC-----------------
Amphiura-5      MNEIEEDC-----------------
Amphiura-10     REEIAETC-----------------
Amphiura-11     REEIAETC-----------------
Amphiura-4      KGEVQELCP----------------
Amphiura-14     KHEIQELC-----------------
Ophioderma-10   MEELDETC-----------------
Ophioderma-3    MEEVNETC-----------------
Ophioderma-8    MEEVNETC-----------------
Amphiura-13     WEEIKETC-----------------
Amphiura-17     VEEINEVC-----------------
Amphiura-18     LEEVNEYC-----------------
Ophioderma-1    VEEIDEVC-----------------
Ophioderma-2    NEEVNESC-----------------
Ophioderma-4    SEEISEHC-----------------
Ophioderma-7    TEEIREHC-----------------
Amphiura-12     VQEVYDHCDKHFDGSI-YDLFRS--
Acanthaster-5   YEEVYELC-----------------
Patiria-6       YEEVYELC-----------------
Asterias-4      YEEVYELC-----------------
Pisaster-2      YEEVYEVC-----------------
Pisaster-5      VEEVNESC-----------------
Asterias-1      IEEISESC-----------------
Pisaster-3      VEEISESC-----------------
Asterias-2      SEEIHEHCP----------------
Pisaster-4      AEEIHEHCPKVY-------------
Patiria-2       WEEILEYC-----------------
Acanthaster-6   WEEILEYC-----------------
Acanthaster-7   WEEVLEYC-----------------
Patiria-1       WEEMLEHC-----------------
Acanthaster-2   NEEIYEVC-----------------
Acanthaster-1   IEEIYESC-----------------
Acanthaster-8   DEEIYET------------------
Acanthaster-9   DEEIYEIC-----------------
Patiria-5       NREVRKHC-----------------
Patiria-8       NEEVREHC-----------------
Ophioderma-5    LEEIVELSKTMCSSS----------
Ophioderma-6    YEEIEEVC-----------------
Ophioderma-9    YEEIEEVC-----------------
Asterias-3      WEEIEEVC-----------------
Pisaster-1      WEEIEEVC-----------------
Acanthaster-3   WEEIEEVC-----------------
Patiria-7       WEEIEEVC-----------------
Acanthaster-4   WEEVEEQCH----------------
Patiria-3       WEEVEEHCH----------------
Patiria-4       WEEVEEHCH----------------
```

Figure S9-continued.

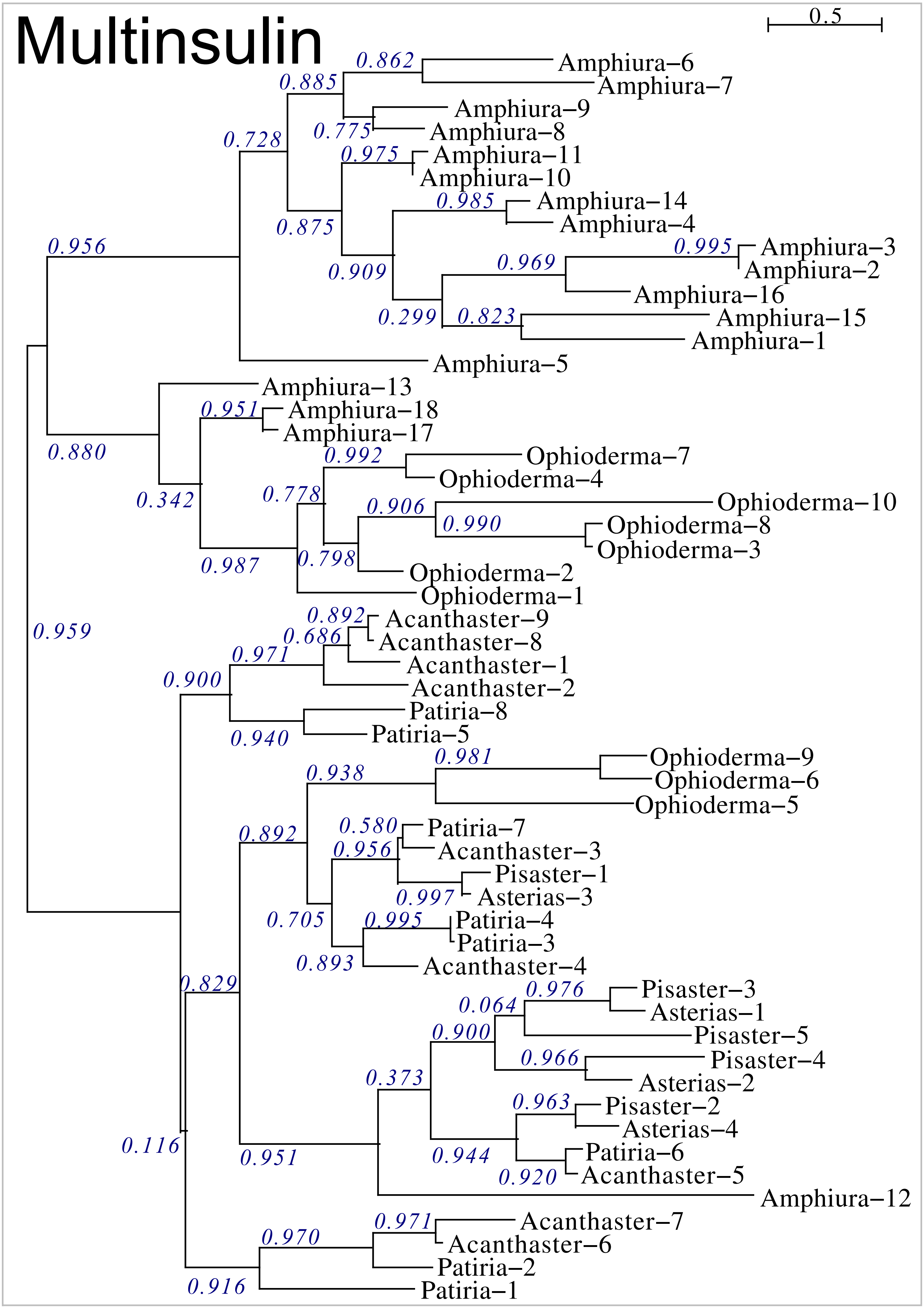


Figure S10. Sequence similarity tree of ambulacrarian multinsulins.

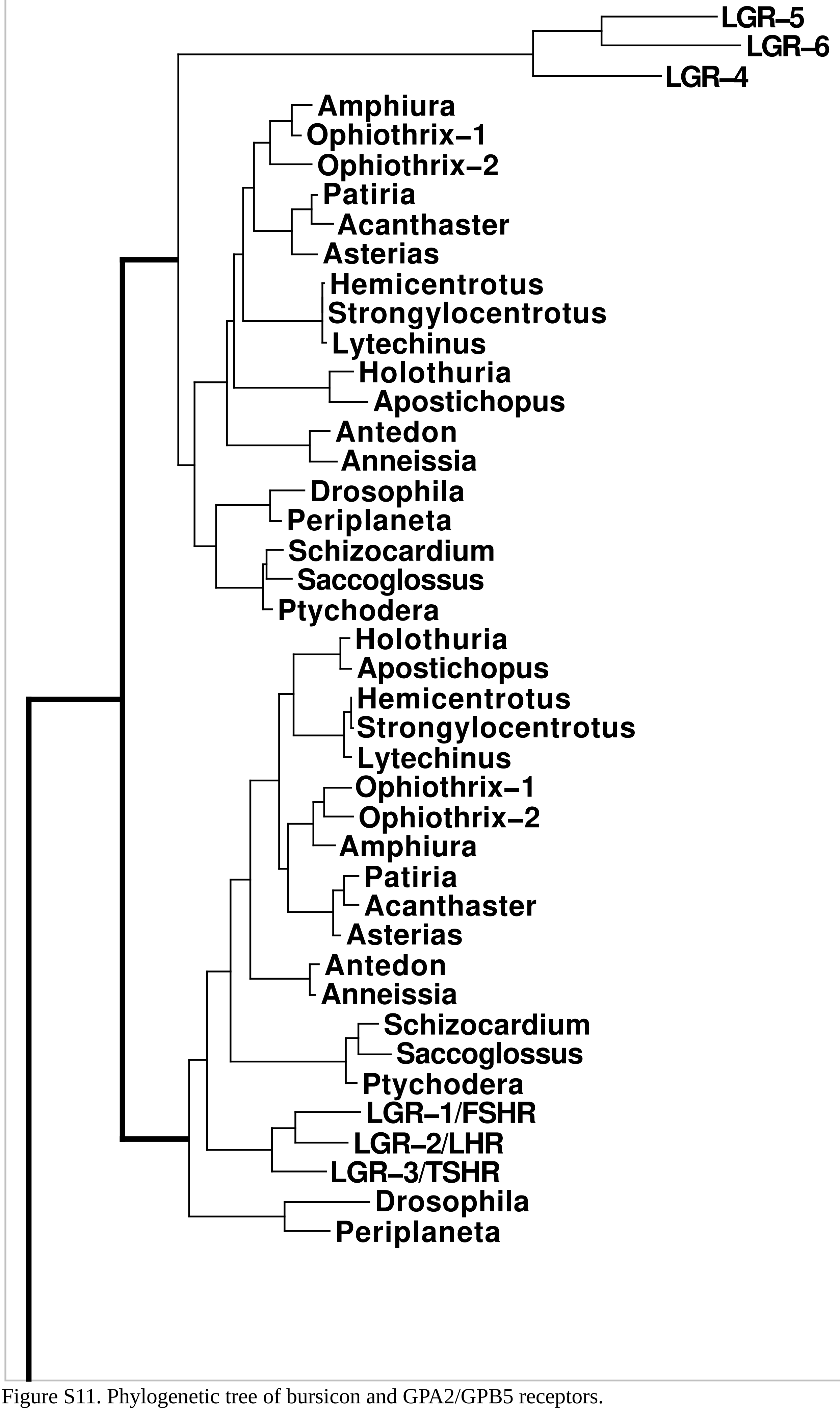




Figure S11. Phylogenetic tree of bursicon and GPA2/GPB5 receptors.

```
Ptychodera          ----------------------------------------------------------
Periplaneta         ----------------------------------------------------------
Anneissia           ----------------------------------------------------------
Apostichopus        ----------------------------------------------------------
Strongylocentrotus  ----------------------------------------------------------
Patiria             MEMKEPAEDGNKTKIRAGSDGTPRPRNILTSRSSRASRSSRGDGFRGSVWSAASSMLH

Ptychodera          ---------------MFIVITLLATLPG-------------------------VTSNE
Periplaneta         ----MQCRRSVACGLAVLAVVLA---------------------------LAALTFLL
Anneissia           -------------------------------------------MCSSKLAGLICIFVL
Apostichopus        ----------------------------------MLTRTGKWILLSVTM----IALTS
Strongylocentrotus  -------------MKMEL-ILRVVTFTSA-L--AAMSRTGSCQ--------------W
Patiria             RLRTRQCKVLVLCLVLLLILVLAVTFIGVSLTGNDVAQPGNNEFIPATVKTEYEELIG

Ptychodera          LPQSFSCPDGYFQCGNV-TECVHQKYFCDGKADCVDGFDEIPTKCRDNFGWANLISRI
Periplaneta         SQTHGACAEGTFPCGNS-SICLPQRFICNGHIDCPNGEDESKYTCTDVHGSNFNQDIV
Anneissia           LPFTYCCQSGTFPCGDDKEQCFPQEFQCNDVVDCTNAADEQ--DCEEKAGFGDDFDRR
Apostichopus        PCWAFECPKGEFVCGNL-TKCVKQKYQCNEIDDCGNNADEEEEVCRNFEGWVKSMDKV
Strongylocentrotus  ANDLIECDPGFFPCGNS-TECVAQQFQCDDIAHCQNAADES--GCAIADGWLAHFDYV
Patiria             FSSEYTCPKGEFRCGNL-THCVAQKYQCNGEDDCGNNADET--ECEHNEGWIKTFDEQ
                    LDLa

Ptychodera          TTFDKSNNDTLQPRNFIRRCELDDYPEACECT-FTSVSCANANLTGIPTNISGSVTHL
Periplaneta         IN----RNY----SQAFNDCQLESLPRGCLCRDKTWVFCRELGLTHVPQDISTNVTRL
Anneissia           HNTTIVLKYD-VGNYNTNVCLIPVYPPVCKCGNKTEVDCSATGMTTIPQAISRNISRL
Apostichopus        F----DEKD-----NIERECELTNYPPVCDCVNGEKLICSNKGLTEVPSVPPNT----
Strongylocentrotus  HPVSVGETK------ALNECALAQYPDGCVCSKLTQLDCSEFNLNEVPPNLPENITRL
Patiria             NPVPVDLDW-----QINQECGLYGFPDVCVCFEETELKCIQVNLTQVPQSVSANLTRL

Ptychodera          YVSGNKIKRLEPGIFKEYPNLLTLSLFDNAVEKLEEDTFQGLPRLEKLYLNDNKIKEI
Periplaneta         ILSNN-SITLSQDSFKKYHNLELIHLGWNQISDIPNGMFKGQRTLRRLFLSKNSLTEL
Anneissia           TLLGNDIQTILNNSFVNFEQLHKLTLAQNGLVRMEINAFNGLTELKQLYLSENNIKEI
Apostichopus        ---------------------SHLILMNNQIHDIPLTAFEGLTNLKVLYLSENPISTL
Strongylocentrotus  YLKSNNIVNIGSELLANYSSLVYLNLMGNRIHEIAENSFRNLRNLKRLFLSENNISRI
Patiria             NLNGN-MVHLEDGAFERYTKLGSLNLMQNGIRELPRDVFRGLTDLDKLYLSTNRISTI
                    LRR

Ptychodera          EDGAFRSLQKLAWLMLNNNHLKSLGEGMFEGLESVYYLNLKDNEVTEADFGTLTFTHM
Periplaneta         RPGVLQDLPHLSWLVLQDNCISDIDLDTFQYL-NLTWLDMTRNNLT--LKGK-IFPDL
Anneissia           PSGVFLPLKRLEWLVLDYNELTTLLGNPFKGLLSLYWLSVRHNKINDANIRT-VLENL
Apostichopus        RAGVFRDLINLEWLFLVNNTLRSLDDDAFIGLRKVYWLRLNSNLLEDLSDGT-PFRHM
Strongylocentrotus  QTGAFNGLSGLHWLFLNDNKMRTLEPGIFLGLEGLYWLDMRNNEIRDLQEGE-ALKDL
Patiria             TPGTFRFLQNLTWLFLNENEIETLDEDVFEGLHKVYWLMLQKNKIGDLNHGV-AFRDL
                    LRR

Ptychodera          PSLEWLNLGDNDIKHLHTGMF--PNNSALKVLFLGSNLIQSIHEDTLRGLDTLEDLDL
Periplaneta         PELKQLILEDNTIETIGEFTL--TNLKALQALDLTKNRIRYIHPKAFANLWNLEELEL
Anneissia           PSLAWLELDHNKITRIEAEDF--ARDTPMSTLSLKNNEVSYVHPLAFTSLINLLDLDL
Apostichopus        QELKWCDLGSNKITKITSGTF--QGETGLQTLDLSNNKIHTISKGAFLGQDNLTVLNL
Strongylocentrotus  KSLVWLDFEGNDIRSINARSF--AGNTSLTVLILRNNSIESIERDAFTSLGTLVELDL
Patiria             PKLMWLDMSNNPLHHLSADNFSASGNPPLSILMLSNCNISTIQKDTFQAFRELTALHV
                    LRR

Ptychodera          SDNLLSTIPAKTFHELISLAKLNLCNNRKLRRLPLELFSNLHQLRTLNLTGVKITNID
Periplaneta         SGNQLVSLSSSVFYNLTFLMKLSLSNN-PFRLLPTQLFRRLENLRSLGLVDTDIQNID
Anneissia           CGNKISELSPMQFKNLTKLDRLALGFN-LFDSLPLNIFDDLVNLQSLRLEGITINNID
Apostichopus        TLNNLSSVPKGLSSELPMLTTLSLSFN-PIQ-LDELSFSGFHNLKEIDLVGVDIPNIN
Strongylocentrotus  SHNQISHLPEGIFDGLHTLEILRLRNN-PLFSLPVDVFRGMKKLVDLDLAEIIIGNIN
Patiria             SGNKIGHFPSGLFRNMANLTVLTIANN-PATSLPEDLFDDLVLLDDLNLGGLVINNIS
                    LRR

Ptychodera          TKMFNSMQKLEHLYFAKFHYCSYASHVRDCSPKSDGISSLENLLANKVLRCFIWIVAA
Periplaneta         VELFAHCPKLEFLYFSKFYYCTYAPAVRICRPNTDGVSSLKHLLVKPALRVTVWVVAA
Anneissia           QRMFEKMEKLSIVYFSSFQYCSYAPHVRNCNPKTDGISSIYNLLESTVLRVSVWTIAT
Apostichopus        NAMFEQINSLTILHFERFEYCKFAHHVRNCNPKTDGISSTENLLEGAILRISVWTIAL
Strongylocentrotus  VDMFQGLTRLEKVAFDKFEYCRYAPHVRTCTPRSDGISSFENLLKNVVLRVFVWTIGI
Patiria             ARMFKGMTNLQHIEFFKFEYCRYAAHVRTCKPRSDGISSFENLLKDGILRVSVWTIAL
                    Hinge region
```

Figure S12. Sequence alignment of various gonadulin receptors.

```
Ptychodera          TTLVCNLAVILGRMCIMKIENKVHSLLIKNLCGADLLMGIYLVIIGSHDLMYRNHYYI
Periplaneta         VTCAGNALVLWGRF-SSRDENRVHSLVIRNLAVSDLLMGVYLLVIGTQDCQFRDRYHK
Anneissia           MTFIGNFGVFISRL-LMKAEHRVYSLFIMNLCAADFLMSIYLFTIAAQDYRFRDVYNQ
Apostichopus        LTTCGNLNVFISRI-AIHAENKVHSIVVSNLCAADFMMGLYLFAIGIQDARFRNNYID
Strongylocentrotus  ITSVGNLGVLVSRA-FMKAENKVHTVVVINLCTADFLMGIYLLIIGTHDVKYRDNYNQ
Patiria             LCFLGNIGVLISRY-LMKAENRIHSLVVVNLCTADFCMSIYLFVIGYHDAKFRNQFNT
                        TM1                            TM2

Ptychodera          YSHKWMTSWTCSLCGFLAVFSSETTLLVLTVISVQRFLCIAFQTRMIH-FKYSYGVIS
Periplaneta         EAHSWMTSWKCTVVGMVAMTSSEVSVLILVFLSMERFLLIAVPFGGHQNMSMRTTCLS
Anneissia           NARQWVESTGCKMSGFLAMVSSEVSVLLLALISVERFLCIVFPYEFHR-LKVKEAAVI
Apostichopus        YALDWSESWGCHFVGFLALLSTEASVLTLTFISLERFFIVAFPYRFHR-LNIKEVVSV
Strongylocentrotus  HALEWTNGITCKVSGLLAMISSEVSVLTLMFISLERYFIIVYPYSFQR-IKTRRAIMV
Patiria             HALEWMQGSACKFAGFMAMFSSEVSVFMLTFISLERFICIVYPYRLHR-LTSKEALVI
                                      TM3

Ptychodera          IAAAWITAAALAAFPLTM-SSFNHFYGKNGACFPLLDPNDPFEAGWQYSVAMFLGLNA
Periplaneta         LAVIWFIGVCLAAVPVMYWINSTRFYGTNGMCFPL-HIDDPFLLGWEYSAFIFLGVNF
Anneissia           ISSIWAVGILIAWIPLLAEGYFGNFYGTNGVCFPL-HIHGPRIKGWEYSVAIFLGMNL
Apostichopus        LILIWIVAFVLAASPLIFTEYFKNFYGANGVCFPL-HIHDPRLEGWEYSIFIFLGINT
Strongylocentrotus  LAFIWFFGTLLAVVPIIPVDYFGNFYGSNGVCFPL-HLHEPRMRGWEYSVVVFLGINS
Patiria             MTIIWFLGALVAWIPLISVGYFVDFYGSNGVCFPL-HIHDPWLQGWEYSAFIFLGLNA
                        TM4

Ptychodera          VCFCLIVCSYVGMFYSIRETRNAAFKDINLRRRGDMNFAKRFVFIVLSNALCLVPIML
Periplaneta         SSLLLIAVLYAWMFVSIWRTRHATP----ISSVGDFEFVIRFFFIVLTDAGCWAPIFA
Anneissia           LALILIIVSYIGMFISIRTTRKAAV---QIGYKGDMTYARRFFFIVLTDSLCWIPIII
Apostichopus        FSFMVIIFSYCGMFQSIQKTRKRTT---T-LLPGDINYAKRFFFIVLTDCICWLPLAV
Strongylocentrotus  TAFVAIALSYTGMFLSIRKTRRAAA---QCGMKGDMAYAKRFFFVILTDSLCWLPIAV
Patiria             TCFTAIAISYTGMFISIQQTRKAAT---HLGRRGDMNYAKRFLFVVMTDALCWLPIAI
                     TM5                                              TM6

Ptychodera          ITAMVFTDIGVPGQVYAWTTVFVVPINSASDPLLYTLTTVPCKQGCLGMLDKKLKRQK
Periplaneta         LKLVALGRVHIPADLYAWVVVFILPVNSAVNPLLYTFTTPKYRGQLCQLG----CPPQ
Anneissia           LKVISLCNIPIGDNLYAWIVIFVLPLNSALNPILYTLSTTTFSQWFLKLL----KIHK
Apostichopus        VKVLAVADMPVSESIYAWVVIFILPINSALNPILYTLSTGPFRRVLLQKI----KNRG
Strongylocentrotus  LKVMSLCNFIISETLYGWIIVFVLPINSALNPILYTLSTTSFAMWFMRQL----KRGK
Patiria             LKILSLCSYEIPATLYGWIVIFVLPINSALNPILYTLSTTSFTQWFQKHL----KRRK
                                           TM7

Ptychodera          ESGTRKMLVSSTTGSRSSRDSSGTTRLTSSGS-----SGSPSQDT-------------
Periplaneta         TLATKRHSSADTELSRSSTNRNAVDRRADN----GLYAMLPARDPAHT----------
Anneissia           QVS---HNYVTYSPQRTARRKSGIAEENGSTDGMMSYANRPSLQTNCS----------
Apostichopus        KHT---ETGAIPKNERFSNGTN---DTAGETLRLQ--SNLTM-----MGGNKHEANK-
Strongylocentrotus  FKT---SGCYIGSNVDMSELRG---RIEESGA-FTSYTNMPKRASFFLPTRKHHVSSK
Patiria             GKS---SNGSLGSKNEFSTTRG---RIGSTTDQPTEYSSIPKRSYDSNNS--------

Ptychodera          ---KPCIEM----DTAQPLVQN--------SSVGQIQSGQS----------------C
Periplaneta         ----------------------------------------------------------
Anneissia           ----DLLSTAISEDAAEDVRNNTQYKAQHSD--------------NEQPAST-YFVTC
Apostichopus        -----T----MAENTC------------------------------------------
Strongylocentrotus  VEAIPAVSSSVPSRSATTTAMTNNIRPLHTINEGEIVDNHSQSQGNHQLRRTPIVRLC
Patiria             ---SPVLEEE----QCSTV---------------------------------------

Ptychodera          A---
Periplaneta         ----
Anneissia           V---
Apostichopus        ----
Strongylocentrotus  DLME
Patiria             ----
```

Figure S12-continued.

```
Melibe               MEESITPREGSGSEAEESGSTTTATAKLWQHTRARTHYLVFLGVMICVALIVVLVCVT
Drosophila           MSIAI-----------------------------MPHLPITFTLAILLAIASNEGAQG
Periplaneta          ----------------------------------------------------------
Anneissia            MEY-S-----------------------------DKCRPFTKVLHCLLVLQLFFSFHL
Lytechinus           ---------------------------------------MNLALLMVVVTSFFDTSLA
Strongylocentrotus   ---------------------------------------MNWAVLMVVVSSFFGTPQV
Apostichopus         ----M-----------------------------KTMALICHKILQVLIAALFLGV--
Holothuria           ---------------------------------------MMTNILVFLIISMVTRD--
Acanthaster          ------------------------------------MRKLHNALRMLFVLHLCA----
Patiria              ------------------------------------MHKLRNALRMLLILHVCA----
Saccoglossus         ----------------------------------MNLIGDVRKLKYLNTALR-A----
Ptychodera           ---------------------------------------MKKSLRFLLILSL-V----
Shizocardium         ----------------------------------------------------------

Melibe               ST-------------------------TEK-----------------AEALTNALGAS
Drosophila           VESATRTAIEAIRTGIGTKPETEIADATEAEAPVREVISLLGIIDGAESDILVPDADD
Periplaneta          -------------------------------------MFLLQVLF-AVITVTETVGTF
Anneissia            VL--TC-------------------RVI-------------INTCPDVQEVTSKSPHG
Lytechinus           SH-NTK-------------------DDHLDDA--------P-VIRHRRQAPMSTDPHA
Strongylocentrotus   SN-ATK-------------------GDDLDDA--------PSLIRYRRQSPMTSDPHA
Apostichopus         ---TFA-------------------GKEVDQS--------VELHRVRRQ--SSSDVHG
Holothuria           ---AVA-------------------TQGDLDV--------YGARRVRRQ--SNSDNHG
Acanthaster          ----SR-------------------RAEQDGA--------IIRHIQR----RAVDAHK
Patiria              ----AR-------------------RVKQDSA---------LRHLQR----RTVDAHK
Saccoglossus         ----VH-------------------GYSQRQR--------MGRVHQR----IV--RGT
Ptychodera           ----SS-------------------SVSKRGR--------LGHSPHR----RLPRSFF
Shizocardium         ----------------------------------------------------------

Melibe               RCPSNTFRCPSDGRCINRTLVCDGRKQCTDGADEGTVCD--KDKMD-------LWSDM
Drosophila           KCPGGYFHCNTTAQCVPQRANCDGSVDCDDASDEVN---CVNEVD--AKYWDHLYRKQ
Periplaneta          LCKEGQFKCNKSRLCVPQDRNCDGIPDCPDKSDEWD---CSDKLG--SQFWDHLYRKN
Anneissia            DCPGFRFPCQNSSQCVPGNFICNGISDCDNGSDELP-EECATNNPLYTNIWTHIYGSE
Lytechinus           HCGE-WFPCINSTQCIPQEAICDGNIDCDNESDESD-EECIDDKN--RRVYTFMDQGK
Strongylocentrotus   HCGE-WFPCINSTQCVPQEAICNGIFDCDNKMDESD-EECIDDIN--RRIYTFKDQGK
Apostichopus         HCQVTEFPCGNSTECVLQSKHCDSVVDCANGADEDE-EDCDDMDN--TNNFDFLFGKK
Holothuria           HCGL-DFVCGDSMQCVTQSQQCDGVADCDNGADEEE-EECTDEDD--YRHFDEIIGRK
Acanthaster          HCGK-DFPCLNSTQCVPQDSICDGTPDCDNGSDEWEVEECRDWNL--AKMWDNFFGTK
Patiria              HCGV-DFPCLNSTQCVPQDAICDGTPDCDNGSDEWEMEECRDLNL--AKMWDDFFGTK
Saccoglossus         CSGI-QVKCTNSTQCIFRDQLCDDHVDCDNGSDEWS-SECTDEDD--NTNWDILIGQK
Ptychodera           CLDN-SFKCLNSTDCIFREYLCDGIEDCDNGSDEWD-EECTDYND--GENWDIKIGDK
Shizocardium         ----------------------------------------------------------
                     [                    LDLa                 ]

Melibe               FNKRLDADR-------------------------------------------------
Drosophila           PFGRHDNLR-------------------------------------------------
Periplaneta          PSAELDEED-------------------------------------------------
Anneissia            RRGGDGDSSDED----------------------------------------------
Lytechinus           TDDSSEEDD-------------------------------------------------
Strongylocentrotus   QDDSSEEED-------------------------------------------------
Apostichopus         ED-S--EER-------------------------------------------------
Holothuria           EE-L--DEL-------------------------------------------------
Acanthaster          NDDSGEEEEDDLDQGLRPQ---QTVFDAHGHCGEEFPCMNSTQCVPREAICNGKPDCD
Patiria              D-ESGEEEEDDVNQGLSPDPLPQRSFDAHGHCGEDFPCINSTQCVPQEAICNGEPDCD
Saccoglossus         SDSEEEED--------------------------------------------------
Ptychodera           DDESSEEEGEDD----------------------------------------------
Shizocardium         ----------------------------------------------------------
                                                 [                 LDLa    

Melibe               --------------------------------EKEAKNCVWDKIPSRCVCR-------
Drosophila           -----------------------------------IGECLWPNENFSCPC--------
Periplaneta          -----------------------------------L-VCELMINSSECAC--------
Anneissia            --------------------------------FNPDNVCENGTYPPNCTCYLKCISRH
Lytechinus           ---------------------------------FIPPACDIGTYPEECRCVLLYEDPM
Strongylocentrotus   ---------------------------------FIPPACDAGTYPDVCRCVLLHEDPM
Apostichopus         ---------------------------------PPPVTCTDTIFPDECECFVVFDEES
Holothuria           ---------------------------------PPREPCNGTVHPEECKCFVVAEE--
Acanthaster          NGSDEEEIV-CRNKKM----------RDNIEDIRQEIPCEEGTFPESCDCFVEIEQTR
Patiria              NGSDEWELEECRDWNLAKMWDDFFGDKAEDEDVRREIPCEKGTFPESCDCFVEIEETR
Saccoglossus         -------------------------------NESETIECDNGSYPEECNCTHFILREN
Ptychodera           -------------------------------LGGSRIPCAKDTYPNVCNCTYIEFENG
Shizocardium         ----------------------------------------------------------
                     [    ]
```

Figure S13. Sequence alignment of dilp7 receptors.

```
Melibe              ----------------------------------------------EKGKLYC---RD
Drosophila          ----------------------------------------------RGDEILC---RF
Periplaneta         ----------------------------------------------VERSVLC---NN
Anneissia           GGN-------------------A-------------------DDGECESEVFCYTENT
Lytechinus          AHF------DYYES----MLQDA--YP-----SVFAEPPIGATSEPLGLEIAC---TS
Strongylocentrotus  VNF------DYYES-----LQDV--YP-----SAFADPLFGESGEPMGLEIAC---TS
Apostichopus        ---------------------------------------GLENDEPIGLQLDC---QL
Holothuria          -------------------------------------------GEIIGLELLC---PQ
Acanthaster         TAELPSLASPYNTTGDASTYRDA--YDIDSNSTDGVRYGSEVRGVVIGIRLDC---RA
Patiria             -KDLPSLASPFNATGDASMGAGVDDVGTETNVTDGIPYGLGVQGVVVGVRLDC---SA
Saccoglossus        GFM----------TSE----------------N--------QFYTIIDTRLDC---SH
Ptychodera          HNA----------TSR----------------SDE----HPNDNSPYAIHIDC---SD
Shizocardium        ----------------------------------------------------------

Melibe              QDLSAIAHPLPQYT-SVLELSGN-ITSISRLSIGRLPSLHTIIVIHSRVKEIGALTFD
Drosophila          QQLTDIPERLPQHDLATLDLTGNNFETIHETFFSELPDVDSLVLKFCSIREIASHAFD
Periplaneta         MGLTLPPPLLPARGIELLDLTGSSFPVLNATIMDTLPDLEILVLSHAGIKELQQEAFS
Anneissia           TGFTTVPRDLPTNT-VLLDLSDNYISVVRKGDLDHLPYLRTLYFSRNRNLRFESDALT
Lytechinus          KRLERVPEGLPPGT-LYMDLSDNRITHLHGDSFQNLTRLRELVITHNQMQSMDEDVFY
Strongylocentrotus  KQLDRVPEALPADT-LYLDLSDNRITHLHDDSFKNLTMLRELVITHNQMQTMDEDVFY
Apostichopus        SELESVPDNLPRDT-VYLNLAGNSLSNITVETFKTLTKLRRLDLQSNRIGSLHPDSFR
Holothuria          SGLTSVPRGLPRET-IYLNLAANEITTIDKDSFDEVTSLKRLYLQSNKISILHPQSLN
Acanthaster         KQLTNVPRNLPNNT-ISLDLSDNKITHLTKKDLTNLPQLRQLLLSRNKLKRMEEGAFK
Patiria             RQLTEFPRNLPNKT-ISLDLSDNKITSLTKKDMEHLTHLRQLLLSRNKLQNVEEGTFE
Saccoglossus        RNLTSIPGGIPPNI-TKLNLRYNSIQEVKAGDLYGLYDLTELLLSRNIINDIAERALD
Ptychodera          RNLTAVPQNIPDNT-TEIDLSQNSIETIRKADFAGLFMVKNLLLSKNSIREIEEGSFD
Shizocardium        -----------------------MIQEIGSEDLGGLVMLRELLLKKNNLTKLNSSSLS
                    [========================= LRR ==========================]

Melibe              EC--------------------------------------------------------
Drosophila          RLAD------------------------------------------------------
Periplaneta         KA--------------------------------------------------------
Anneissia           ----------------------------------------------------------
Lytechinus          DTR------------------------------------------------SLEMLDL
Strongylocentrotus  DTT------------------------------------------------SLEMLDL
Apostichopus        QLR------------------------------------------------SLELLDL
Holothuria          KLL------------------------------------------------KLEILDF
Acanthaster         PLT-------------------------------------------------------
Patiria             SLT-------------------------------------------------------
Saccoglossus        HLANLMVLDLEANKLKTLPESVFSFLNNLVDLDLSKNDLGTLPIYIFKNLNKLQKMDL
Ptychodera          DLRDLLILSLDRNKLTSLPASVFQTVTNLRELSLELNDISNIPDTVFQNLGHLEELDL
Shizocardium        DLVNLEKLAIESNKLTTIPVDIFSKMKNVQILTLAKNDLQHLPVGVFASLRRLEELDL
                    [========================= LRR ==========================]

Melibe              -----------------------------------------PSLANVHFTFNKLSRFS
Drosophila          -----------------------------------------NPLRTLYMDDNKLPHLP
Periplaneta         -----------------------------------------PSLHTLYLDNNQLRTLP
Anneissia           ---------------------------------------NNLTISELNLVGCQMGTIP
Lytechinus          SDNRIERITSAHFDSLRRLSLLRLSRNH-FRCEDGDCFRNLVSLRELELVACGITELT
Strongylocentrotus  SDNLIEIITAAHFDNLQKLTLLRLSRNH-FRCEGGDCFRNLVSLRELELVACGITELT
Apostichopus        SNNRISSLSKEYFLFNGILKELLLKHNPIQRLEDAELFINLQMLETLDFFGAQISELR
Holothuria          TNNKISWLGSQHLQQLFSLKELTLKHNPIRRLNDSDMFRHMYSLQTLNFLGAQISELR
Acanthaster         ------------------------------------------DLETLRMIACDLEDVQ
Patiria             ------------------------------------------DLQTLRMIACDLEDVP
Saccoglossus        SGNEIRLI-QSELFLGQGSLDVLHLGWNRPLEISEHAFSALRNLTRMRLLDNGLSHLR
Ptychodera          SGNEIRLL-GKNLLRNQVTLKKLQINWNRPMEIEFGAFDNTVNLNTLRLIDNGISHIR
Shizocardium        SGNELRRL-DKDLFENQSGLKTLQLNWNKPLELEFGVFDRLKDLQILGLLDTGLSHIR
                    [========================= LRR ==========================]
```

Figure S13-continued.

```
Melibe              SSVFAKRNSIIYLNLMGNAIKELTSGDFRRMKSLKKLDLRDNKITSIDEGVFEDTPHL
Drosophila          EHFFPEGNQLSILILARNHLHHLKRSDFLNLQKLQELDLRGNRIGNFEAEVFARLPNL
Periplaneta         EPALQPGNILDTLILRNNELSKLPKNAFLNLQYLRELDLRHNHLADVDGDVFQPLVRL
Anneissia           DGLFSGMSHLMTLHLQYNNLVTINKDTLKGLDWLMELNLNHNEISDLNPMAFNHTPNL
Lytechinus          PDMFRNLHSLSVLQLSYNAITEIMPSHMNGLEGLQDLYLSSNHISHIENGSFDGMDRL
Strongylocentrotus  PDLFSNLHSLSVLQLSYNAITEIIPSHMNGLERLQDLYLSSNHISNIEHGTFDGMDNL
Apostichopus        TGVFSPLENLRKLVLAHNEIEALTQGNMEGLRNLLELDLQNNEVSFIEDGTFKEMPML
Holothuria          TGIFEPLTKLTRLVLSHNQIEALTQGNMAGLSSLQTLDVQSNDISFIEDGAFIELESL
Acanthaster         QRMFAAQKRLVTLDLRYNKLTRLVKNSLFGLRNVKYFDIRGNQLSEIETGVFEDTPKL
Patiria             PRMFATQNRLVTLDLSYNILTRLVRGSLFGLHNLEYLDIRGNRLAEIETGVFKDSSRI
Saccoglossus        SGMFRNLNRLKELDLERNQILSIKKYDLQGLTDLSHLKLKNNTIRRIEDGAFDHTPSI
Ptychodera          SGIFRNLNKLTKLDLERNFILSIKKYDLQGLHNLSSLQLRNNSLTRIEAGAFDHTVKL
Shizocardium        SGIFCNLGKLTKLDLERNKLLSIKKYDLEGLDNLKDLLLTNNTIQRIESGAFDHVGKL
                                            LRR

Melibe              ISLDMRGNLLTELVPGMF-DGLQDLRKLYLGNNAIAEIAPGAFASLSKLRAINLARNR
Drosophila          EVLYLNENHLKRLDPDRFPRTLLNLHTLSLAYNQIEDIAANTF-PFPRLRYLFLAGNR
Periplaneta         EILYLNENRLDSVPL---SLPLPSLQLLSLTHNHIAWIEPGAFSQLPRLKHLYLSWNQ
Anneissia           VILNIQYNKLQVLSPPLL-EPLTQLKTLGMSFNQISEIKNGFFESQTHLQDLLLMGNK
Lytechinus          LKILLHDNKLKNISVGVF-PFMPELVILKLDENEIDMIEPGSFSLVMNLQHFTLMENK
Strongylocentrotus  LKLLLHDNKLTNISVGIF-PVMPELQILKLDENEIAIIEPGSFALIENVQHFTLMKNR
Apostichopus        NELRLHGNKIHEVKQGMF-EDLANLTILLLDQNLIHTIEPGSFSKLFKLQHITMSNNM
Holothuria          EELRLHGNKLTKVKRGMF-TNLAQLRTLLLDQNEIESIEPGTFAELTNLQHFTISDNL
Acanthaster         YFMLVSQNKLSSIPANLL-RPLRELRTLDVHRNDISVIETGAFSTNTKLIELNLRDNK
Patiria             YFMHISQNRLSSIPAELF-RPLRDLRSLNIHRNDISSIECGSFSTNTKLRDLNLGDNK
Saccoglossus        SELWLSMNDLTEVKRSVF-LPLKNLKTLDLSFNKIEEFEDGALSNSVILAILLLAGNR
Ptychodera          KQLRLSLNTLPAIKKDVF-EPLQELKILDLSHNKIEDIESGSLSNSTRLSHLLLASNK
Shizocardium        ETIGLSFNDLVEIKKDVF-SPLKEIRSLDFSFNKIQRIETGAFANSRLMTHLFISDNN
                                            LRR

Melibe              IRSLRDGGLDGLGNLTTLDLHYNGMHTIESHAFKSLVSARSVDLRSNHLISIGNGTFS
Drosophila          LSHIRDETFCNLSNLQGLHLNENRIEGFDLEAFACLKNLSSLLLTGNRFQTLDSRVLK
Periplaneta         LWVLRNDTFDNLPELLSLSLNGNKIRLLEVNTFKEVPLLTSLYLQDNQLKLLDSRVLR
Anneissia           LTVIKVGMFQNLYGLLSMTLRENNIREVQANSLDHIESLQSLSLAKNPFESLPDHVFC
Lytechinus          LSRIELGMFDGLENVTDMSLRQNEIKAIEENAFDNLIRLDHY-LRENKLTEVPRELFT
Strongylocentrotus  LSRIEVGMFDGLGNVTDMSLLENEIKVIEENAFDNLIKLESLDLQKNKLTEVPRELFT
Apostichopus        LTHLTVGMFDGLSQVSSLSLYANKITEMENGTFEPLIKVSSLNLKDNLFRVVPRGIFD
Holothuria          LTHLPVGMFEGLSRVVSLGLFDNHIETMENGTFAPLLNVESMNLERNPFRIVPRGVFD
Acanthaster         LTEIRRGIFHSLTSTITLSLSNNSIRHLEQDAFAGMNNLQTLKLTKNPFTSLPVGIFD
Patiria             LTEIKRGLFHNLTSVITLSLRNNSIRHFEEDAFAGMNNLQTLKLNQNPFTSVPIGIFD
Saccoglossus        LPFVRTGMFANLTKVSNLPLFDNEIAWFDADSLQDMESLRTLDLQHNPVQRLPMVLFD
Ptychodera          LSILRNGMFTSLSSVTNLDLLDNEIRSLEPGCFDDMLSLYTLDLRENPFTLLPRGLFD
Shizocardium        LHDIRIGMFSGLATVTNMELNNNDIEVVEAGAFDDMTSLYTLDFQKNPFTRLPRGIFD
                                            LRR

Melibe              YMH-QLHYIYFDKFHMCLYAQHSQQCLPLGDGISSKENLLESIVLRICVWVVALLSLG
Drosophila          NLT-SLDYIYFSWFHLCSAAMNVRVCDPHGDGISSKLHLLDNQILRGSVWVMASIAVV
Periplaneta         HLQ-NLEHIYFDEFKLCSAALHVRVCEPKGDGISSLQHLLDSIVLRASVWVMAVLACA
Anneissia           AMTNTLDTIEFDSFSFCGFAPHARHCYPSTDGLSSRDHMLKSVILRISVWIVGSIALF
Lytechinus          PLE-NLNNIYFDYFFMCGYAPNVRVCLPKGDGISNVDDLLGNWLLRGAVWLVALLGSF
Strongylocentrotus  RLK-NLNNIYFDFFFMCGYAPNVRVCLPKGDGISNVDDLLGNWLLRGAVWLVALLGCF
Apostichopus        VMP-MLGHIYFDHFFLCGFAPHVKDCDPKSDGLSTSEDLLGSIVLRVAVWLVSILASF
Holothuria          VMP-KLSHIYFDHFILCGFAPHVKDCNPKGDGVSNSENLLGSIVLRVSVWMVSILATF
Acanthaster         QLI-SLKAIYFDHFSLCGYAPHVRLCMPKSDGISTAENLLGNILLRFAVWFVALLASL
Patiria             KLR-SLKAIYFDHFSLCGYAPHVRLCMPKSDGISTAENLLANNLLRFAVWFVALLASL
Saccoglossus        DLP-SLEWVHFDHFWMCGYAPTVRHCTPRGDGISSVENLLDNVVLRVMVWVVAMLAFV
Ptychodera          NLL-SFEWIYFDHFSLCGYAPTVRHCSPRGDGISSIENLLDNMVLRIMVWVVATLAFI
Shizocardium        KLGESLQWIYFDHFSMCGYAPTIRHCEPRGNGISSIENLLDNMVLRIMVWVVALLAFI
                                                                   TM1

Melibe              GNMVVFLGRF-LLREDNQIHSFFIKNLSMADLLMGLYLLIIAIHDIKFRGQYILHDES
Drosophila          GNLLVLLGRYFYKSRSNVEHSLYLRHLAASDFLMGIYLTLIACADISFRGEYIKYEET
Periplaneta         GNTVVLLGRLVARE-PNPVHSLYIKTLALADLLMGIYLMIIASADWHYRDVYVRHDFQ
Anneissia           GNLFVLTTRLFVNE-TKRVHAFLIMNLAFADLLMGIYLYLIAIHGALFRDNYILYDLV
Lytechinus          GNLLVILARCFVSE-DNKVHSFFIMNLAVADFLMGLYLLIIGLHDVMFRGEYIFKDLN
Strongylocentrotus  GNLMVIFARCFVSE-DNKVHSFFIMNLAVADFLMGLYLLIIGLHDVMFRGEYIFKDLD
Apostichopus        GNLIVLFARCVSKE-DNKVHSFFIINLAISDFLMGAYLFIVAVHDVMYRGVYILHDVD
Holothuria          GNLVVLFARCLSKD-DNKVHSFFIINLAVSDFLMGAYLFIVAIHDVIYRGSYITYDLD
Acanthaster         GNAFVLLARCFVNE-DKKTHSFFIMNLAVADLLMGLYLLIIGIHDVIFRGSYILHDLT
Patiria             GNAFVLLARCFVKE-DKKTHSFFIMNLAVADLLMGLYLLIIGIHDVIFRGVYILHDLA
Saccoglossus        GNLFVLVARCILKE-DNRIHSFFIMNLSLADLLMGLYLFIIALHDVMYRGEYIKFDLE
Ptychodera          GNIFVLIVRCVLKE-DNRVHSFFIVNLSFADLLMGLYLFIIAVHDVIYRGEYIKFDLS
Shizocardium        GNIFVLIARCILKE-DNRVHAFFIMNLSFADLLMGIYLFVIAIHDVIYRGEYIKYDLS
                                                 TM2
```

Figure S13-continued.

```
Melibe              WRNSWECDFAGVLSTLSTEMSVLTLSVITLDRYISIMYPLSLRRGLRVAYFVMALTW
Drosophila          WRHSGVCAFAGFLSTFSCQSSTLLLTLVTWDRLMSVTRPLKPRDTEKVRIVLRLLLW
Periplaneta         WRHSPVCNACGFLSTLSCEASVLILTLITRDRLVSVTRPLDRRQPSLRHATLLVAVLW
Anneissia           WRSSDICTIAGCLSILSSMMSVFTLSVITLDRYVSIVHPFSYESKSLFRAAILMGVLW
Lytechinus          WRSGWVCRMCGLLSLLSSEMSVLTLTVITSDRFISIVHPFKFRQRHLAHAVILMVAFW
Strongylocentrotus  WRSGWVCKMCGLLSLLSSEMSVLTLTVITSDRFISIVHPFKFRQRHLAHAVILMVGFW
Apostichopus        WRSSWVCQLCGAFSLLSSEMSVFTLTIITADRYFCIVHPFRFRSRNLIPAAILMGCLW
Holothuria          WRSGWVCQMCGAFSLLSSEMSVFTLTVITADRYFCIVHPFRFRNRQILPAAILMGVLW
Acanthaster         WRNSSVCKLSGFLSLLSSEVSIMTLTVITLDRFLSIVHPFRFKNRSLVHARLLMVFLW
Patiria             WRTSSVCKLSGFLSLLSSEVSIMTLTVITMDRFLSIVHPFRFKNRSLVHARLLMVLLW
Saccoglossus        WRLGWVCQMCGFLSLVSSEASVLTLAIITMDRFICIVYPFKFKNRNMKIAVFAMLLIW
Ptychodera          WRTSWICKLCGFLSFVSSEASVLILTVITLDRLFSIVYPFKFKNRTIKLAFLIMATMW
Shizocardium        WRLSWVCQLCGFLSLVSSEASVLTLTIITLDRFICIVYPFKFKNRNLKLAFWVMLMVW
                                   TM3

Melibe              TVCIM-LVSLPVMG--LEYFGDSFYRDNAVCTPLHLHHPRAKGWEYSSFLFLGINFCS
Drosophila          GISFGL-AAAPLLP--NPYFGSHFYGNNGVCLSLHIHDPYAKGWEYSALLFILVNTLS
Periplaneta         LVAAVV-AALPLCAETVEYFGEEFYGGNGVCLPLHVQDPFADGWEYSAAMFIALNTVA
Anneissia           LFGITL-VLIPIIM--KQTFGELFYGSNGVCLPLQIRHPWATGWVFTTTLFVGLNSVF
Lytechinus          LAAMLI-AILPVLH--RSYFGEFFYGGNGVCLPLQFDRPFDHGWEFTLVVFVLFNLIA
Strongylocentrotus  LAAMLI-AILPVLH--RNYFGEFFYGGNGVCLPLQFDRPFDHGWEFTLVVFVLFNLFA
Apostichopus        IIGITL-SILPLLH--RSYFGDFFYGANGVCLPLQFDRPFDDGWLFSLFVFVVANLLS
Holothuria          LLGITL-AMLPLVY--KTYFGDFFYGANGVCLPLQFDRPFDDGWVFSLFVFVIANLLS
Acanthaster         LLGIAL-ATIPLLH--TAYFGEFYYGGNGVCLPLQIDQPFADGWEFSLVIFVVFNLVA
Patiria             LLGVAL-ATIPLLH--TEYFGEFYYGGNGVCLPLQIDQPFANGWQFSLVIFVVFNLVA
Saccoglossus        ISCVII-ATLPLID-AITYFGDFFYGGNGVCLPLHIDDPKADGWEYSLVIFVLVNFMA
Ptychodera          SICIFL-AVLPLID-HLEYFSVFFYGGNGVCLPLHIDEPMTNGWEYSLFIFTFINLVA
Shizocardium        LACVFL-AILPLIH-VITYFGAFFYGGNGVCLPLHIDEPKANGWEYSLFIFVLVNFVA
                    TM4                                                     TM

Melibe              FAFIAYAYVAMFYSIHMSALPLRTSRDSKERCLVQRFFFIVLTDFVCWVPIIIIKILA
Drosophila          LIFILFSYIRMLQAIRDSGGGMRSTHSGRENVVATRFAIIVTTDCACWLPIIVVKLAA
Periplaneta         LLFISFAYWRMLRVVRSSGLSLRSTQERQDKAVAQRFVIIVATDCLCWIPIIGVKLAA
Anneissia           FLFVVYAYIAMFVMIRRSD-DVRSTKKSQDSAILVRFTLLVFTDMLCWLPICVIKVLA
Lytechinus          FIFILYAYARSKAIPRMNKMGKRSTKESQDWNLLKRFTIIVATDFICWMPIILVKIAS
Strongylocentrotus  FLFILYAYAQMFATVRKSSLAMRSTKESQDWNLLKRFTIIVATDFICWMPIILVKIAS
Apostichopus        FTFIVWAYIRMFFTIRRSNLAARSTKVSQDYALLKRFTVIVATDFLCWMPIIIVKFVT
Holothuria          FLFIVWAYIRMFTTIHRSNLAARSTKVSQDYALLKRFTVIVATDFLCWMPIIIVKFVT
Acanthaster         FTFISYAYLMMFMTIRRSNLAMRSTKKNQDWALVKRFTLIVATDFVCWMPIIIVKFVA
Patiria             FAFISYAYLMMFVTIRRSNLAMRSTKKNQDWALVKRFTLIVATDFVCWMPIIIVKFVA
Saccoglossus        FLFIVFAYMAMFTSIKKSRSNIRSTKENQDLLLVKRFTLIVATDFICWMPIIIIKFAA
Ptychodera          FLFIAFAYAAMFNAIRQSRTNIRSTTENQDYMLAKRFTLIVATDFVCWMPIIILKLVA
Shizocardium        FVFISFAYVAMFSSIRKSRTNIRSTKENQDLVLVKRFTLIVATDFICWMPIIIVKFAA
                    M5                                        TM6

Melibe              LSGVAITVDLYAWVIVFILPVNSALNPLLYTLTTQYFKKKLFSKFSAVVFRPVLRSD-
Drosophila          LSGCEISPDLYAWLAVLVLPVNSALNPVLYTLTTAAFKQQLRRYCHTLPSCSLVNNET
Periplaneta         LAGARVDQQLCAWLAVLILPVNSALNPVLYTLTTSFFKSQMSRLLHAWKSRDTGPSHD
Anneissia           ISGYPVSGDVYSWLAVFVLPVNAALNPILYTITSKLFRQQLR----IILHKCRRATGY
Lytechinus          YAGIVIPQSVHAWFAIFVLPVNSALNPILYTVTTQFFRQKFLRPIRRICGRQKQGKAY
Strongylocentrotus  YSGIAIPQSVHAWFAIFVLPVNSALNPILYTVTTQFFRQRFLRPIRRLCGRQKQGKTY
Apostichopus        YGGVAIPPAAHAWFAIFILPVNSALNPILYTMTTRQFKRMIFHPSRTFMGKQKSTAGH
Holothuria          YGGVIIPPTAHAWFAIFILPVNSALNPILYTMTTRQFKRMLFHPSRALMGMSKSAGGN
Acanthaster         LGGVSVSQSVYAWFAIFVLPINSALNPILYTMTTVLFRQKILAPLGIVKAKRK--KGY
Patiria             LGGVSVSNSVYAWFAIFVLPINSALNPILYTMTTVLFKQKVLAPLGIVRAKRK--KGY
Saccoglossus        LAGAQISGGVYAWVAVFILPVNSALNPILYTMTTKMFKQKFMKLLGISNKKKN--AG-
Ptychodera          LAGVKVNGDIYAWMAVFILPVNSALNPILYTMTTKLFKQKMLRALGVHLRD----KA-
Shizocardium        FAGAKINGDVYAWIAIFILPVNSALNPILYTLTTRMFKQKMLRAFGVATK-----KE-
                                 TM7
```

Figure S13-continued.

```
Melibe              ----------KDS----N----------------------QSRSSMKNNTLYSRHMHD
Drosophila          RSQ----------TQTAYESGLSVSLAHLGGGVGGGSGRK--RMSHRQMSYL------
Periplaneta         SAG------------SLSNIQLSTRKRHLGSQPPTRSSTL-----RSHSSYKSIRGT-
Anneissia           RRG---RRDDDSDSFKRSGTRLSLI--PVRSKASLKSKSS--IDKTKSNS--------
Lytechinus          QSG---SYDDSITGTKITATKLSII--SHNGKPRAGSVNG--RLNSTKGSHGHHRP--
Strongylocentrotus  QSG---SYDDSITGTKISATKLSVI--SHNGKPRAGSVNG--RLISTKGSHVHHRP--
Apostichopus        EMSQSMSRTRDREIGHTATSRLSVI--SHSNTMKNGT---------------------
Holothuria          DMSQSVSRSRDREMGNTATSRLSVI--SHSHQMKNGGGTA--TPG-------------
Acanthaster         ITG--VSVDETSTMSKNSGTRLSII--SNKS--RGGSLNG--R-----------FNSQ
Patiria             LTG--ISVDDTSTMSKNSGTRLSII--STKS--RGGSVNG--R-----------FSSH
Saccoglossus        --------DESYSASKTTGTRLSSV--ASSR--NRISLNG--NIKIYTYSIPFICSTI
Ptychodera          --------DESLTASKTTGMRVSTF--GQR---SRSTTST--NGKARNGSFHDNVQDN
Shizocardium        --------DDSFTASKTSGTRLSST--GSR---TRISISA--K--NSNGSIGTRATSL

Melibe              LEMDLLRRYGNASRSGT----QTTTTTLSTSR---IPHNGKATSVRPRAQSDNALNIR
Drosophila          ----------------------------------------------------------
Periplaneta         ---------------AV-----------------------------------------
Anneissia           ----------------------------------------------------------
Lytechinus          ---SEVQPWLSPDSSGRPSHLKRPSLRMEDMSCRKE-----------------PEVVV
Strongylocentrotus  ---SEVQPWLTPDSSGHPSHPKRPSVRVEDTSCREE-----------------PEVVV
Apostichopus        -----------------PTTPRNG--HMENLSHED-----------------------
Holothuria          ---------------GTPAAERNA--HMENMSNDD-----------------------
Acanthaster         KKLKNLSSLDSTDESVTCSAAQTT---SLKIKKHRA-----------------A-TAD
Patiria             KKLKNFDSLDSSDESVICTATKTT---T----KKRA-----------------V-TVE
Saccoglossus        FRLKTILFFIIKKSDTPAS--STT---LRDIKLAAA-----------------PKSLR
Ptychodera          -QFDRDVAFDNQGADMS----------GEPIALSVQ-----------------AEHLR
Shizocardium        RTTSREQTYPVKDCNYV----------ELTTIYTPP-----------------PE---

Melibe              PNVGNA---SQPISNI-----------------
Drosophila          ---------------------------------
Periplaneta         ---------------------------------
Anneissia           ---------------------------------
Lytechinus          VHSNSQN--RIECSDVAEPLIET----------
Strongylocentrotus  VHSNSQN--RIESSDVVEPLIET----------
Apostichopus        ---NTNN--EDRDQNMDSPE-------------
Holothuria          ---NKKE--RNEE--MAAL--------------
Acanthaster         YHELPTSDP-DCAPSGVNDDME-----------
Patiria             YHQLPVSDS-DERPLASTVT-------------
Saccoglossus        RTLIEQGDGSSEVRSAAAIEIQRAWRGYWARWS
Ptychodera          GTE------------------------------
Shizocardium        ---------------------------------
```

Figure S13-continued.

```
Apostichopus        ----MMRILQSVVFLALLLLM------------------VVNISQSSPPDC-EETDIF
Anneissia           ------MEH---RLVPMLLAIAWISLC-----------VVSKSELHGSVSC-KSTDVK
Strongylocentrotus  ---MMMG-RQAFIFLVLGIA------------------LVDIVAGVNLQQCDVEVEFE
Patiria             ------MR-LTSILLVVFLISLFLA------------SDARLRNKDETTSCENEADFE
Ptychodera          ------MGRYLGSCHRFLLYVIWFS--------------VTVAETQHQVVCDIDTELE
Periplaneta         MLFQLLPNKASLVFVICLQIVYICSSTNEPVLGSGVMHGAVPGGSVVMPCNSSEAVFH
Lymnaea             ------MA-TMSGTTIVCL--IYLT----TMLGNSQGVNLKIESPSPPTLCSVEGTFH

Apostichopus        CDN---KCLNVTMQCNNVYDCVGGEDESFLICGCRHDEFTCSD-GCIPLHERCDLNCS
Anneissia           CGT--GECVAANSSCNSISDCPDGSDESPAYCGCRHDEFQCYN-SCISLFGRCDGTCD
Strongylocentrotus  CEP--GVCIPRNLTCNGIFDCELGQDEEVAFCGCAKDEFPCSH-SCIPIELRCDTNCD
Patiria             CEP--GVCIDASLACNSAYDCESMADETVMLCGCPSGEFRCLN-SCIPLEKRCDRHCD
Ptychodera          CSP--GVCIDLQYRCDGIPDCENGHDESTLVCGCPSHEFQCNETTCIDLIRRCDWSMD
Periplaneta         CDA--VVCVPKSSLCNGIPDCQDGRDESVAQCGCLPNEFRCSN-SCVDLVRRCDRQPD
Lymnaea             CDDGMLQCVLMGSKCDGVSDCENGMDESVETCGCLQSEFQCNHTTCIDKILRCDRNDD
                                             LDLa

Apostichopus        CSS-CEDEDECEGYSCPPTHN-KCK-NHYCIPQINFCDFIDDCGDNSDEENCNHPPCS
Anneissia           CDF-CEDERDCDLHQCPSTHM-KCATSHICIHLSLQCNFVDDCGDGSDE-MCEHRECY
Strongylocentrotus  CEVDCADMDGCEAFPCLEGYE-KCTKNHYCIAKHLWCNFVDDCGDNSDEDLCYHPDCW
Patiria             CQDDCADEDHCDSFQCLPGFQ-KCE-NHFCMPTYAWCDFVDDCGDMSDEALCERLDCW
Ptychodera          CPD-GSDEDNCETFQCLHTHS-KCQNTHKCIPRDKTCDFIDDCGDKSDEEDCYRRPCY
Periplaneta         CAD-GEDEQDCEAYVCPT-THFKC-NNHFCIPGDSVCNFVDDCGDGSDENKCSHRKCW
Lymnaea             CSN-GLDERECDIYICPLGTHVKC-DNHFCVPRDKQCDFLDDCGDNSDEKICERRECV
                                             LDLa

Apostichopus        SMFLFECENHQCISNAYACDGIEDCFDGSDER--HCEEEN-FHICEDGNKIHEGRKCN
Anneissia           T-HEFQCANGECLKPGYRCNGEVNCRGGDTSDEDNCIENYDYIMCR-SGRIFRNLFCD
Strongylocentrotus  K-GEFQCSNKQCINTWFVCDGSQDCIDGSDE--ARCGEEH-FVLCENGKKVYEHEWCD
Patiria             Y-GQFQCINQQCINSFFLCDGIQDCTDGTDEMESTCSPGN-FADCGDGTRVRAQQFCN
Ptychodera          AYGEFRCENEQCISSYIVCDGINDCLDGSDEV--QCQQED-FVTCSDGTQIHRHQWCD
Periplaneta         N-LEFKCNNGECIRPGFVCDGTQDCKDGSDET--DCGPDQ-FALCGDGSRVHRYYWCD
Lymnaea             A-TEFKCNNSQCVAFGNLCDGLVDCVDGSDEDQVACDSDK-YFQCAEGSLIKKEFVCD
                                             LDLa

Apostichopus        MVRDCPSGS-DEHGCEC--QVYQTQCPAEKSCFDNDIHCNSFCECAD----------C
Anneissia           FFIDCELTEEDEKQNCSCDSADGFMCS-TGVCILRSKLCDGQCDCRDL---------C
Strongylocentrotus  RLVDCPDNEADETNCVC--TADEYRCP-NGKCLRPSVRCNGVCDCLS----------C
Patiria             GDAKCQLNKAVEINCDC--SDDEFRCG-NGKCLPPAKQCDRLCDCSN----------C
Ptychodera          GIPDCRESHDDEVNCYC--AEDDFRCG-NGRCISRAYLCDLSCDCAGT---------C
Periplaneta         GWPDCPDNHADELKCQNCSGVDEFLCP-NGRCIRRANVCDSQCDCVPPPGSAAVNVTC
Lymnaea             GWVDCKLTFADELNCKLCD-EDDFRCS-DTRCIQKSNVCDGYCDCKT----------C
                                             LDLa

Apostichopus        SDETNCDEI---------CDV-SNGIYCPWMNYYIGSYRLSGCVKTERFCDGTPDCLG
Anneissia           EDENNCGDT--------VCSA-GSDYFCA---------HSFRCLNSEFVCNGYNDCEN
Strongylocentrotus  DDEKECADEDM-------CNYETNGILCQ-----IANNKPSRCVRKEYICDGFGDCPN
Patiria             EDEEGCDNKDD-------CDTDT-HIFCP-----PVDGKPPVCLQKQFICDGRRDCPI
Ptychodera          EDEKNCSSG--------LCTL-ENHFWCD---------EFSRCLHKDTICDGKFDCFD
Periplaneta         DDEVDCDLYYTNINGIHLCTQ-TETLNCIEP---NQNRDKDRCIHREFICDRTNDCQN
Lymnaea             DDEEVCANN---TYG---CPM-DTKYMC------RSIYGEPRCIDKDNVCNMINDCRD
                                             LDLa

Apostichopus        Y---QDEMFCSVPLHVP------------VRQC---PSTRDPRYIPES-EFCDN--SS
Anneissia           KDDE----LCESRYKTD----VGYECKLNSDMCQL---DNNAFCYPTE-IFCER--RP
Strongylocentrotus  K--AD-EFNCQL--PGNTTGHCDSEGNPPYMMCDIKDGADDPRCLPTLKVQCDRYRRP
Patiria             F--GWDEDDCQTLPDGESVASCNVPG---TVSC----GAGDPRCPPSKYVKCDK--RF
Ptychodera          ---WSDEYGCGDDVNCS---THEYENQMEFFRCG-GVESDDGRCIPER-AVCDY--QR
Periplaneta         GNYLSDEFGCPDDLDVI----------GESIVC------SDNRSLPKA-LECDF--KW
Lymnaea             GNVGTDEYYCSNDSECK---N--FQAAMGFFYC------PEERCLAKH-LYCDL--HP
                                             LDLa

Apostichopus        YCFQGSEETNCTHGERVCSVGQFQCADS--SCVEESLRCNGFWDCKDFSDEIDCPNYI
Anneissia           QCQGGSNGSHCL-VEALCNSNEFKCTNG--MCIPDYLRCDNIADCLDHSDELNCESHP
Strongylocentrotus  HCMYGFDEETCNATRPCDPEVEYECPYG--RCIDLTSRCDAQLDCFDFSDEANCESFE
Patiria             DCLTGVDEQNCTV-MPCPPGQHQCRRGG--YCVKEELWCDGYVDCFDGSDEDNCELFE
Ptychodera          HCILGTDESDERCGHPACMEDEFQCNNGHVKCIPIVQRCDTNPDCLDGSDERNCEDYL
Periplaneta         DCLEGNDEHG-CV-PDACAEGEYTCKNG--QCVNSTQRCDLQFDCWDKSDELNCVNEK
Lymnaea             DCINGEDEQS-CLAPPKCSQDEFQCHHG--KCIPISKRCDSVHDCVDWSDEMNCENHQ
                                             LDLa
```

Figure S14. Sequence alignment GRL101 receptors.

```
Apostichopus        CPNENMFKCRGTGQCLASHQRCNGEVECTYINGT-KEDSDEENCV--LRDCEENEFRC
Anneissia           CEE-GMWKCR-EGHCIEMHMRCDRYVDCANFGQY-RDASDEENCD--FESCKQNEFQC
Strongylocentrotus  CLP-GTWKCH-SGQCIPEKQKCDYTPDCFYVNGT-VDSSDEDKCDYKIFGCREGEFRC
Patiria             CPP-GLRKCA-AGQCISQRLFCDYHLDCFYYKNSTRDVSDEAHCT--RRQCREDEFQC
Ptychodera          CDE-GQLQCNESGQCIHRDMLCNFIKNC--------ADNSDEENCGHVIRNCTEDEFQC
Periplaneta         CPS-GKKKCVHSGQCVSAAFWCDFIVDC--------PDGSDETDCGEKPPLCGPNQFRC
Lymnaea             CAA-NMKSCL-SGHCIEEHKWCNFHREC--------PDGSDEKDCDPR-PVCEANQFRC
                    LDLa

Apostichopus        HSGHCIENWRRCYKAPDEHLYGCKDRSHLWDCGQWECADESQFKCPRDHCISTMSRCN
Anneissia           NNGRCIDESKKCFRDY--SSEECRDNSHLRECEDFECPA-GYYKCQKSHCIPDFRVCD
Strongylocentrotus  HSGQCIPQDEVCFFDST-MKKGCKDRSHLNDCANRTCRE-NEFKCRNAHCVNMSDVCN
Patiria             RSGQCIPASRRCYRDQHDHVQGCADLSHLRNCSDFVCPA-GAFKCARSHCINGTMVCD
Ptychodera          RNGQCIDLSRRCYFDELDDVYGCKDKSHLIGCNDFECSA-DTLKCRNGHCVSQSLVCN
Periplaneta         NNSQCIDASLRCFN-SGNPRYGCADGSHLIGCKNWTCPS-NAFKCRNGPCLNSSMVCD
Lymnaea             KNGQCIDPLQVCV--KGDKYDGCADQSHLINCSQHICLE-GQFRCRKSFCINQTKVCD
                    LDLa

Apostichopus        GLLDCVNNE-DEA------NCLMYCHPA----KYPGCFCKGKIFNCSYTGLTQVNIDHF
Anneissia           VHVDCLLFYDDEV------NCYGKCGTDFEFYSIPDCICNDKTVDCNDLKYGTLDFKEL
Strongylocentrotus  GAVDCLPYWTDED------FCPHECGP-----A-LMCICNHTQMSCINAGLESFHNFYV
Patiria             KNVDCLESWDDEAPRHGDTCIQECAS-------SGCGCSQFDMTCTNIGLNESSFSLN
Ptychodera          GNIDCGLIWTDEE------GCSHICSN-----L--VCMCQDDEMNCEDKNLHEIPYF-I
Periplaneta         HNIDCPGTWEDED------GCLFACSN-----VEPRCECRDIHVNCTNLGLVRVPPD-I
Lymnaea             GTVDCLQGMWDEN------NCRYWCPH-----GQAICQCEGVTMDCTGQKLKEMPVQQM
                    LDLa

Apostichopus        A--DDEVTWFYFNHNQLTV--SALTFGNSTVQNIVHLDLSHNNLQDLPAGLFSMMIGL
Anneissia           ----KDMNIMKFSGT----ELVLESSTFSNVTAITFLDISRNNLMDIGDGYFKDMFFL
Strongylocentrotus  ---EEQISEINLSGNRIQ----ISNEVLSGLPGLLRLDLSNNSLNDIGNDTFKGLANL
Patiria             AKDTRLIIHFSLPGNNLHH-LLQDPTALEGLTRTILLDLSDNGVTFINKAALLNVPNL
Ptychodera          ---EPGYIKLKLGHNYLGEDPNLSNYTFAKLGRLTYLGLEDNNITDLKSFTFSNLWRL
Periplaneta         ---EEEITWFHMANNLINS--TFSEDTFEQLERLLYLDLRNNSISSIPPLAFRNLWRL
Lymnaea             ---EEDLSKLMIGDNLLNL--TSTTFSATYYDKVTYLDLSRNHLTEIPIYSFQNMWKL
                    LRR

Apostichopus        HYLNLGYNGINKLEAQVFSGLTNLKQLVLTGNVIQIIEEGSFLGLTSLKVLDLSHQRL
Anneissia           QTLILSENKITTLKKGAFQGLSNLRYLNLTKNIIENIEADAFDGLDSLTTLDLSYQNI
Strongylocentrotus  RYLNLENNNLRVIRKQTFNGLEGLQTLRLGGNNIHAIEPHAFEGLRNITTLDLSNQNL
Patiria             QTLILADNKITKLGNQTFSSLQHLRTLILRGNAMQTVAGQAFLGLGELTTLDLSQQKI
Ptychodera          LTLNLEDNFLTTLRRDTFDGLGNVRSLYLQGNSIGEIEAHAFKGLSALPTLDLHGQRL
Periplaneta         LTLNLQYNCIRILSNSSFYGLTNLSNLHLQGNGIQVIEAMAFYGLSSLILLDLSWQRI
Lymnaea             THLNLADNNITSLKNGSLLGLSNLKQLHINGNKIETIEEDTFSSMIHLTVLDLSNQRL
                    LRR

Apostichopus        IHIGGNPFVGLYHLTLLNISNNNLQTVEQGIFLKLDNVHTIDMSHNQINFIHENTFNG
Anneissia           QRLPINAFYGLRSLKYLTLRGNKLIFIVGGTFNGLPNLRSLDISENTINKIDGRAFNG
Strongylocentrotus  TEIPNAAFVGLYRLHFLDLSANHIVMVPDGAFYGLYQLKKLDISENAIEVVSRR-TFN
Patiria             QSINKGAFDGLNKLAFLNLSHNRMTDIPRGTFFGLNELKELDISNNKLSQIHAEAFYG
Ptychodera          QWLHINAFAGLSRLLSLNLSHNQIQYIDNGVFNGLSKLLVLDISHNVVIEIDSKVFYG
Periplaneta         HNISQGAFVGLRSLVGLDLSHNEITYLVDGSFNGMPHLLSLDLSSNTIKVISSNVFRS
Lymnaea             THVYKNMFKGLKQITVLNISRNQINSIDNGAFNNLANVRLIDLSGNVIKDIGQKVFMG
                    LRR

Apostichopus        LTNLEKIISDELRFCCVAKH---VKECLPEADEFSSCEDLMRNTFLRFNLWVVGLVAT
Anneissia           LPSLDYLYTDEFRFCCMSEG---VPNCYPEPDAFSSCDDLMSNYVLRVSIWVLGVTAF
Strongylocentrotus  LRLLEELITDEYRFCCMARH---VSKCLPLPDEFSSCEDLMSNPILRMCIWIIGMIAL
Patiria             LPKLRVLHTDEFRFCCMVREMVSVETCTPPPDEFSSCEDLMANVFLRGSIWVIGIIAS
Ptychodera          LMELTELYTDEYRFCCMAKF---VQKCHPPPDQFSSCQDLMSNVVLRGGIWLLGVIAS
Periplaneta         AASLTKLVTDEFRFCCLARH---VKHCLPPPDEFSSCEDLMSNLVLRICVWVLGVLAT
Lymnaea             LPRLVELKTDSYRFCCLAPE---GVKCSPKQDEFSSCEDLMSNHVLRVSIWVLGVIAL
                                  Hinge region                     TM1

Apostichopus        FCNAIVIVTRSGKRRDSRVQSFLITNLAIGDMCMGIYLLIIAIADLNYRGRYAAYESL
Anneissia           LGNLLVIIWRVTHNRENKVHSFLITNLAAGDFCMGVYLLIIAIVDLTYRGEYIIHDME
Strongylocentrotus  IGNLVVIMWRVNSKRDNKVHSFLITNLAVGDMLMGIYLLIIAGVDAYYRGDYIVHDKT
Patiria             IGNLAVIMMRMNNKRDNRVHSFLITNLAIGDFCMGVYLLIIASVDAFYRGYYIIYDAQ
Ptychodera          VGNLVVVCFRLRDRRDNKVHSFLITNLAVGDLCMGVYLIIIAVADTYYRGNYIIHDKV
Periplaneta         FGNILVIGWRMRFKHTNQVHSFLITNLAIGDMFMGSYLLIIAGVDARYRGVYFIHDSD
Lymnaea             VGNFVVIFWRVRDFRGGKVHSFLITNLAIGDFLMGVYLLIIATADTYYRGVYISHDEN
                                  TM2
```

Figure S14-continued.

```
Apostichopus        WKRSLFCKFAGFLSTFSSELSVLSLTFITLHRVSSIVFPFRTKDIGFSRIVWIMTSTW
Anneissia           WRSSSWCRFAGFLSTFSSELSVFSLTIITLQRLTSILFPFRSKNMELGWAMKMMALTW
Strongylocentrotus  WRNSGLCKFAGFLSTFSSELSVFSLTIITLHRLSSIVFPFRIKDMEFTRAVWVMCVSW
Patiria             WRDSALCRFAGFLSTFSSELSVFSLTIITLHRLLSILFPFRIKDMEYSGAVRVMVVTW
Ptychodera          WRQSQFCKIAGFLSTFSSELSVFTLTVITVDRLVCIVFPFRFNRLEFKGAARLMAALW
Periplaneta         WRSSELCHLAGFISTFSSELSVYTLTVITLDRFLVIIFPFRVRRLEMPKTRLLMALGW
Lymnaea             WKQSGLCQFAGFVSTFSSELSVLTLSTITLDRLICILFPLRRTRLGLRQAIIVMSCIW
                                        TM3

Apostichopus        AIVFFLAGIPLLGFSYFDNFYGRSGVCLALHITPDKPNGWEYAVFVFLALNFFSFTFI
Anneissia           LLAIFLAILPLCRVNYFGNFYGRSGVCLALHITGEKTSGWEYAVFIFLAINLLSFIII
Strongylocentrotus  GLVAFLAALPLCGVAYFGNFYGRSGVCLALHITPDKPSGWEYSVFIFLGLNFVSFITI
Patiria             LMVIFLSAFPLFGLSYFGNFYGRSGVCLALHITPDLPKGWEYSVFIFLALNFISFLTI
Ptychodera          LIVFFISAVPLFGLSYFTDYYGRSGVCLALHITNERPDGWEYSVFVFLIINLVNFLII
Periplaneta         LGAAFLSGLPLLQIDYFRNFYGRSGVCLALHITPEKPSGWEYSVFVFLFLNLISFSII
Lymnaea             VLVFLLAVLPLLGFSYFENFYGRSGVCLALHVTPDRRPGWEYSVGVFILLNLLSFVLI
                     TM4                                               TM5

Apostichopus        AVAYMVMFYVARRTQQAAKRSLRGNKGSDAMARKMTLIVFTDFCCWMPIIILGVASLC
Anneissia           ILSYVIMFVVARRTQRAV---R-PQNRGDSMAMRMTVIVLTDFCCWMPIIFLGIASLL
Strongylocentrotus  MVSYGVMFNVARKTQKAAMRSL-KSKGSDSMARRMSVIVFTDFCCWVPIILLGLASLS
Patiria             AVSYSVMFVVARRTQKAVNRSR-DTNTGDAMARRMTLIVMTDFVCWVPIILLGVASLG
Ptychodera          LLSYIAMFIVASKTQQAV-RNR-DLKTESAMAKRITVIVMSDFFCWVPIILLGLASLG
Periplaneta         ALGYLWMYVVARTTQQAV-KKE-QRPSDNAMARRMTLIVATDAACWMPIILLGVLSLG
Lymnaea             ASSYLWMFSVAKKTRSAV-RTA-ESKNDNAMARRMTLIVMTDFCCWVPIIVLGFVSLA
                                                              TM6

Apostichopus        GANIHPPVFAWVAVFVLPLNSAVNPLLYTLWTAPYARKIINSARSTFGRS--TATTEY
Anneissia           GASVSPKVYAWIAVFVLPLNAAINPLLYTLWTSPYVRRFLKKARPSMRSLST-YYTDT
Strongylocentrotus  GAYVPTSVYAWVAVFVMPVNSAVNPILYTLGSAITVRRAIQRFSINPSTT---VTSEY
Patiria             GAKIPPQVYAWVAVFVLPVNSAINPMLYTLLTAPYVRRVMSRARTSLNLSLSTVTTDM
Ptychodera          GAVIPPQVYAWIAVFVLPLNSALNPVLYTLSTPTFVKRTKKMADSIGESFRSRWRTDG
Periplaneta         GITVPPQVFAWVAVFVLPLNAAVNPVLYTISTAPFLGPARRGLTTFKRSCKLSLTMDQ
Lymnaea             GARADDQVYAWIAVFVLPLNSATNPVIYTLSTAPFLGNVRKRANRFRKSFIHSFTGDT
                                   TM7

Apostichopus        KLSANLSTS-------EKVSSTANGRRAQRGL--------ASTSN---------SK--
Anneissia           RHGSVREH--------HRSGNN---FKMRISQ--RSK-------------KTSLKFIP
Strongylocentrotus  KTVMN----------NS-SDRKQGGYKGRP--M-RTMSNTANENCKPA----TTTTTK
Patiria             KQVNH----------GDRQGRMCNGRKGWQKGI-RLPKADANKNAIKLNTIISQSDTE
Ptychodera          RTNSFTVSESQA-----RSS----------------------FSYTGIDRLDRCASVA
Periplaneta         RRTYSSTLGSTFANNCSRCELDYNLYPRLEQTNGKLQEGAGGCHYDGVEDLDSEQDMK
Lymnaea             KHSYVDDGTT--H---SYCEKK-SPYRQLELK--------------------RLRSLN

Apostichopus        -LPPEKSTGSIRS---SDVINDPEEGIRLHDVSHDKGNVYVNPNNNDD------RLGN
Anneissia           SSPPKNNNSTTNKRTDAGVITNDEQYNKLLSDQ-RNKDSYE-----------------
Strongylocentrotus  PKPPPAKEETIKM---IDLGK------RRHDNGDDGEK------------RALTSAGN
Patiria             PDEPATAQTRSEE---LNLLQ------DHHSPTQGAES------------GKEAEAGM
Ptychodera          SSNSATVAIRLRS------FS------GGHNGGADLTGCYDKRTGSDPSTRERCDSGK
Periplaneta         NEAPLFS------------LT------KRHQPHRWRSGSYHATNNVVPPSSSVSTPG-
Lymnaea             SSPPMYY------------NT------ELHSDS-------------------------

Apostichopus        HGNKDAYEKML-----SDKSMDIDGGHSSEE--------------G------------
Anneissia           ----------------------------------------------------------
Strongylocentrotus  NNEK------------------------------------------------------
Patiria             HT--------------------------------------------------------
Ptychodera          SVTFASDSRTWLRRALQAPPKDIKIVEKLREVIIIEFKPEEQSVGAMIIGFLELFSEA
Periplaneta         ------ATILWQSSSCRHGSSSIDTAVSA---------------HGEIIPLRRLHSDS
Lymnaea             ----------------------------------------------------------

Apostichopus        ----------------------------------------------------------
Anneissia           ----------------------------------------------------------
Strongylocentrotus  --------------------TSNNV---------------------------------
Patiria             ----------------------GNV---------------------------------
Ptychodera          QEKHFTRLRDMLKKLSSNDTAHENIIRMLWSGSVSDLLSSQAITEQFPGEFKWCICVE
Periplaneta         -------------------RGNGNK---------------------------------
Lymnaea             ----------------------------------------------------------
```

Figure S14-continued.

```
Apostichopus         ----------------------------------------------------------
Anneissia            ----------------------------------------------------------
Strongylocentrotus   ----------------------------------------------------------
Patiria              ----------------------------------------------------------
Ptychodera           YIEGVTLKTFAKDKIDTRELSRIVIQIAKALDHLRKHKIVYNNLSTSSIIIQRVGQES
Periplaneta          ----------------------------------------------------------
Lymnaea              ----------------------------------------------------------

Apostichopus         ----------------------------------------------------------
Anneissia            ----------------------------------------------------------
Strongylocentrotus   ----------------------------------------------------------
Patiria              ----------------------------------------------------------
Ptychodera           SRKVRPILFDFCKAVDLSVALSTDSEDVDHMNDVMGIARMLNDLCTTVMVTTDSEDTD
Periplaneta          ----------------------------------------------------------
Lymnaea              ----------------------------------------------------------

Apostichopus         ----------------------------------------------------------
Anneissia            ----------------------------------------------------------
Strongylocentrotus   ----------------------------------------------------------
Patiria              ----------------------------------------------------------
Ptychodera           YERTPLFPMSQSVKRNGLRADEVLRMCITECATEPPSAFDILNQLTSLMSDPDIVYYV
Periplaneta          ----------------------------------------------------------
Lymnaea              ----------------------------------------------------------
```

Figure S14-continued.
